# A computational toolbox and step-by-step tutorial for the analysis of neuronal population dynamics in calcium imaging data

**DOI:** 10.1101/103879

**Authors:** Sebastián A. Romano, Verónica Pérez-Schuster, Adrien Jouary, Alessia Candeo, Jonathan Boulanger-Weill, Germán Sumbre

## Abstract

The development of new imaging and optogenetics techniques to study the dynamics of large neuronal circuits is generating datasets of unprecedented volume and complexity, demanding the development of appropriate analysis tools. We present a tutorial for the use of a comprehensive computational toolbox for the analysis of neuronal population activity imaging. It consists of tools for image pre-processing and segmentation, estimation of significant single-neuron single-trial signals, mapping event-related neuronal responses, detection of activity-correlated neuronal clusters, exploration of population dynamics, and analysis of clusters’ features against surrogate control datasets. They are integrated in a modular and versatile processing pipeline, adaptable to different needs. The clustering module is capable of detecting flexible, dynamically activated neuronal assemblies, consistent with the distributed population coding of the brain. We demonstrate the suitability of the toolbox for a variety of calcium imaging datasets, and provide a case study to explain its implementation.

## INTRODUCTION

Brain function relies on the interaction of large neuronal populations. Anomalies of these complex neuronal circuits are associated with diverse brain disorders^1^. Therefore, to understand brain function both in health and disease, it is necessary to explore the activity dynamics of neuronal networks. Recent advances in optical methods and optogenetics provide unprecedented possibilities for functional imaging of large neuronal populations^2–5^, and even whole brains^6,7^, with high spatial (e.g., single-neuron) resolution. These imaging datasets can be analyzed to reveal the responses of spatially distributed neuronal populations to sensory, motor, or task variables^5,8–11^.

Nevertheless, the study of large neuronal circuits imposes new challenges to the processing and analysis of the complex, high-dimensional datasets typically acquired. A popular analysis approach is based on statistical methods of dimensionality reduction^12^. These methods project the high-dimensional data into a low-dimensional space that preserves or reveals underlying features of the data. Remarkably, they have been applied to expose organizing principles of neuronal networks dynamics^13–15^ and to detect neuronal assemblies^9,16–18^. The identification of neuronal assemblies (i.e., neuronal subsets that show correlated activity) is a significant step towards a systemic understanding of neuronal circuits, since they can reflect functional processing modules^2^. Importantly, neuronal assemblies are not fixed, rigid structures with unique functions. On the contrary, they are dynamic, multifunctional, adaptive and overlapping units^3^. For instance, a neuronal assembly could perform a computation at a given time, but at a different time point, a subgroup of this assembly could be part of a different assembly with a different functional role. Hence, neurons could belong to multiple assemblies (i.e., non-exclusive assemblies) and play diverse roles in different brain processes. Indeed, it is in neuronal assemblies’ flexible and distributed nature wherein lies one of the major difficulties in identifying them^4^. The ability to simultaneously record activity in progressively larger neuronal populations is thus paramount in these efforts.

We recently developed a computational framework that applies dimensionality reduction and clustering techniques for the analysis of calcium imaging datasets, which outperforms other traditional algorithms in the effective detection of neuronal assemblies embedded within large neuronal networks^9^. Remarkably, this method is able to detect non-exclusive assemblies that are engaged and disengaged on a moment-to-moment basis, compatible with the distributed and dynamic nature of brain processing^19^. Here, we present a detailed tutorial for the utilization of this framework, implemented in an integrated computational toolbox. The toolbox consists of a complete data processing pipeline designed for the comprehensive analysis of fluorescence imaging data. It includes modules for video pre-processing, morphological image segmentation into regions of interest (ROIs) corresponding to single neurons, extraction of fluorescence signals, analysis of ROI responses to stimulus and/or behavioral variables, detection of assemblies of ROIs, exploratory analysis of network dynamics and the automatic generation of surrogate shuffled datasets to act as controls for statistical purposes. It thus accomplishes a complete computational workflow from raw imaging data to interpretable results on neuronal population dynamics. Typically, the full protocol can be completed on a workstation computer in 1–2 hours, depending on the size of the dataset.

Analytical demands vary greatly depending on the scientific questions and the nature of the datasets. Therefore, the pipeline is organized in flexible sub-modules that can be bypassed, replaced or used in a stand-alone manner, by allowing the user to import and integrate data and/or results from other preferred methods at different critical points of the pipeline.

## DESCRIPTION OF THE TOOLBOX

### Overview

The toolbox includes four main modules (Fig. 1). The first pre-processing module (steps 1-30) contains sub-modules for smooth video registration, automatic detection and interactive manual curation of motion artifacts and morphological single-neuron ROIs, and automatic detection of ROIs’ significant fluorescence events associated with neuronal calcium transients. Since the sub-module defines ROIs that correspond to single neurons, we interchangeably use the terms “ROI” and “neuron” throughout the text. The second module (steps 31-37) allows the characterization of neuronal responses (i.e., tuning curves) with respect to an experimental variable. It also enables mapping of the spatial topography of these responses, setting appropriate color mappings to efficiently visualize the response features across the imaged optical plane. The third module (steps 38-47) performs the detection of neuronal assemblies through different methods, including that which we recently introduced^9^. The fourth module (steps 48-52) is intended for exploratory analysis of these assemblies. It contains sub-modules for interactive exploration of the assemblies’ spatio-temporal organization, and visualization of assemblies’ activity in relation to experimental contexts and/or events (e.g., sensory stimulation, behavioral events, etc.). The fifth and final module (steps 53-55) allows for the creation of surrogate control assemblies, useful for statistical comparison against the original assemblies’ features (spatial, functional, etc; see Box 1). Except for the processing pipeline depicted in Figs. 1, 3 and Supplementary Figs. 1-2, all the other figures presented here are direct screenshots and montages of images and graphs automatically produced by the toolbox.

**Figure 1.**
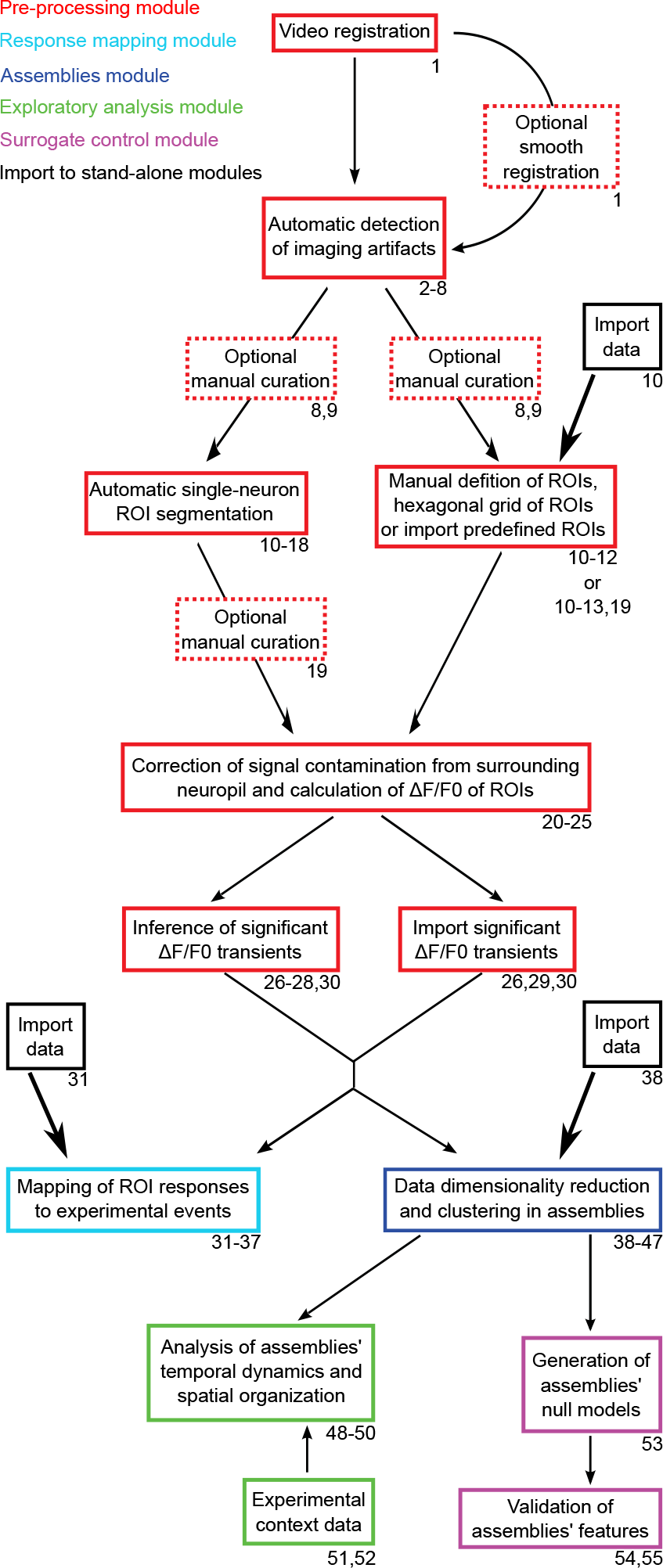
Overview of the toolbox workflow. Colored boxes indicate the processing modules (color code in top left corner). Dashed boxes correspond to optional procedures. Tutorial step numbers related to each procedure are indicated in the bottom-right corner of each box. Single-headed thin and thick arrows respectively depict the processing-pipeline flow, and the option to import pre-analyzed data from other methods into the stand-alone modules of the pipeline.

The methods implemented in this protocol have been extensively tested with *in vivo* imaging data from zebrafish larvae^7,9,20^ and mice (see Fig. 2) expressing genetically encoded calcium indicators (GCaMPs; see Figs. 2b-d and 4-9) or injected calcium indicator dyes (OGB 1-AM; see Fig. 2a). The analyzed imaging data was obtained through two-photon laser scanning fluorescence microscopy^9,10^ (see Figs. 2a-c, 4 and 6-9), and single-photon light-sheet fluorescence microscopy (see Figs. 2d and 5).

**Figure 2.**
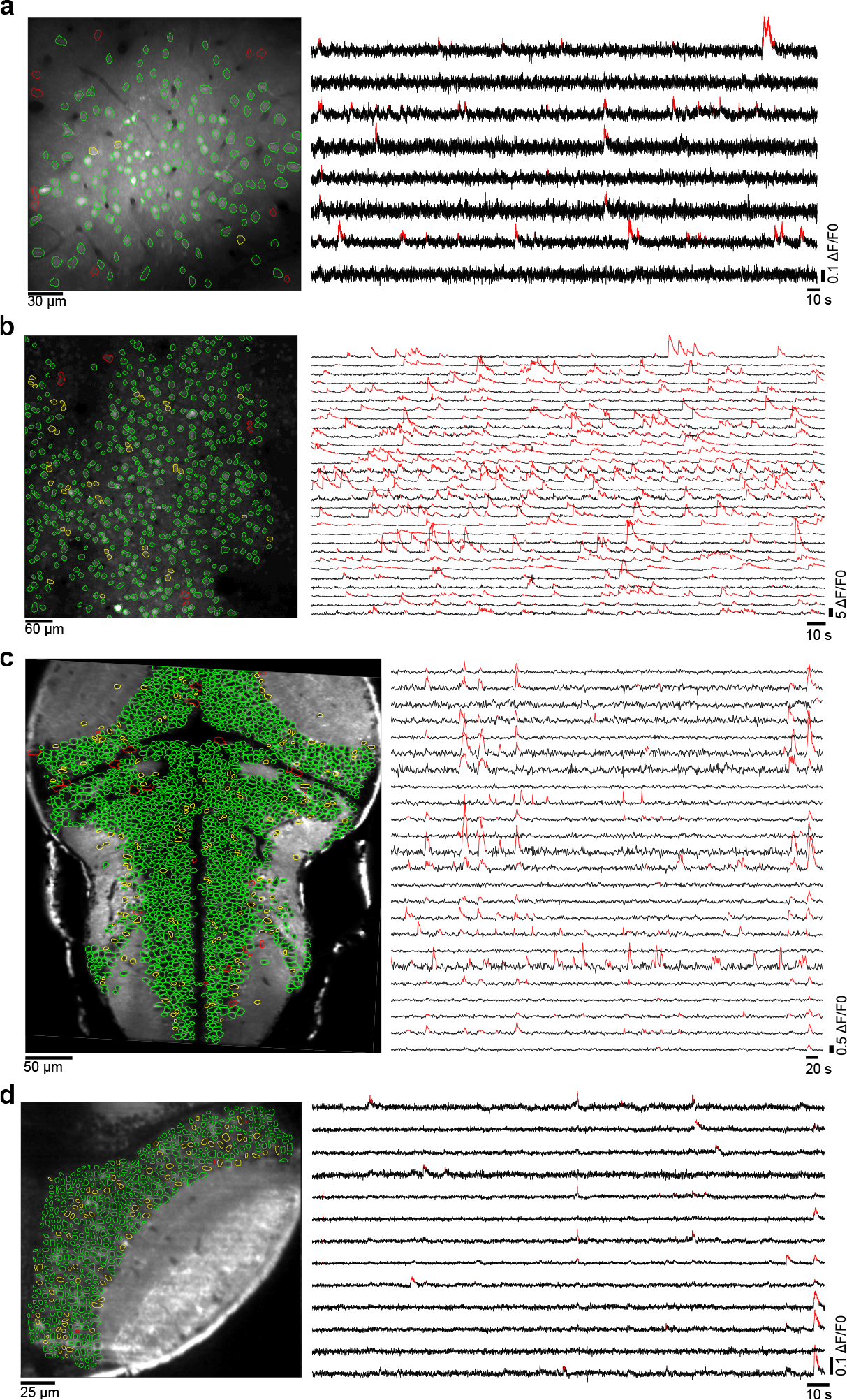
Detection of ROIs corresponding to single neurons. Four examples from different animal models, brain regions, imaging techniques and calcium indicators. Left: imaged optical planes. Perimeters of correct and incorrect automatically detected ROIs are shown in green and red, respectively. Yellow perimeters, detected ROIs that had to be manually curated, or undetected but manually drawn. Right: representative *ΔF/F0* traces (black) and significant fluorescent transients detected (red). **(a)** Mouse primary visual cortex bolus injected with OGB-1 AM (two-photon imaging, 256x256 pixels, 30 Hz sampling rate). Data from Scholl *et al*^21^. **(b)** Mouse somatosensory cortex, where nuclei of excitatory neurons are transgenically labeled with mCherry (two-photon imaging, 256x256 pixels, 7 Hz sampling rate). Calcium dynamics monitored with GCaMP6s. Data from Peron *et al*^11,22^. **(c)** Transgenic zebrafish larva pan-neuronally expressing GCaMP3 (two-photon imaging, 512×256 pixels, 1 Hz sampling rate). **(d)** Right hemisphere of the optic tectum of a transgenic zebrafish larva pan-neuronally expressing GCaMP5 (single-photon light-sheet imaging, 232×242 pixels, 100 Hz sampling rate).

**Figure 3.**
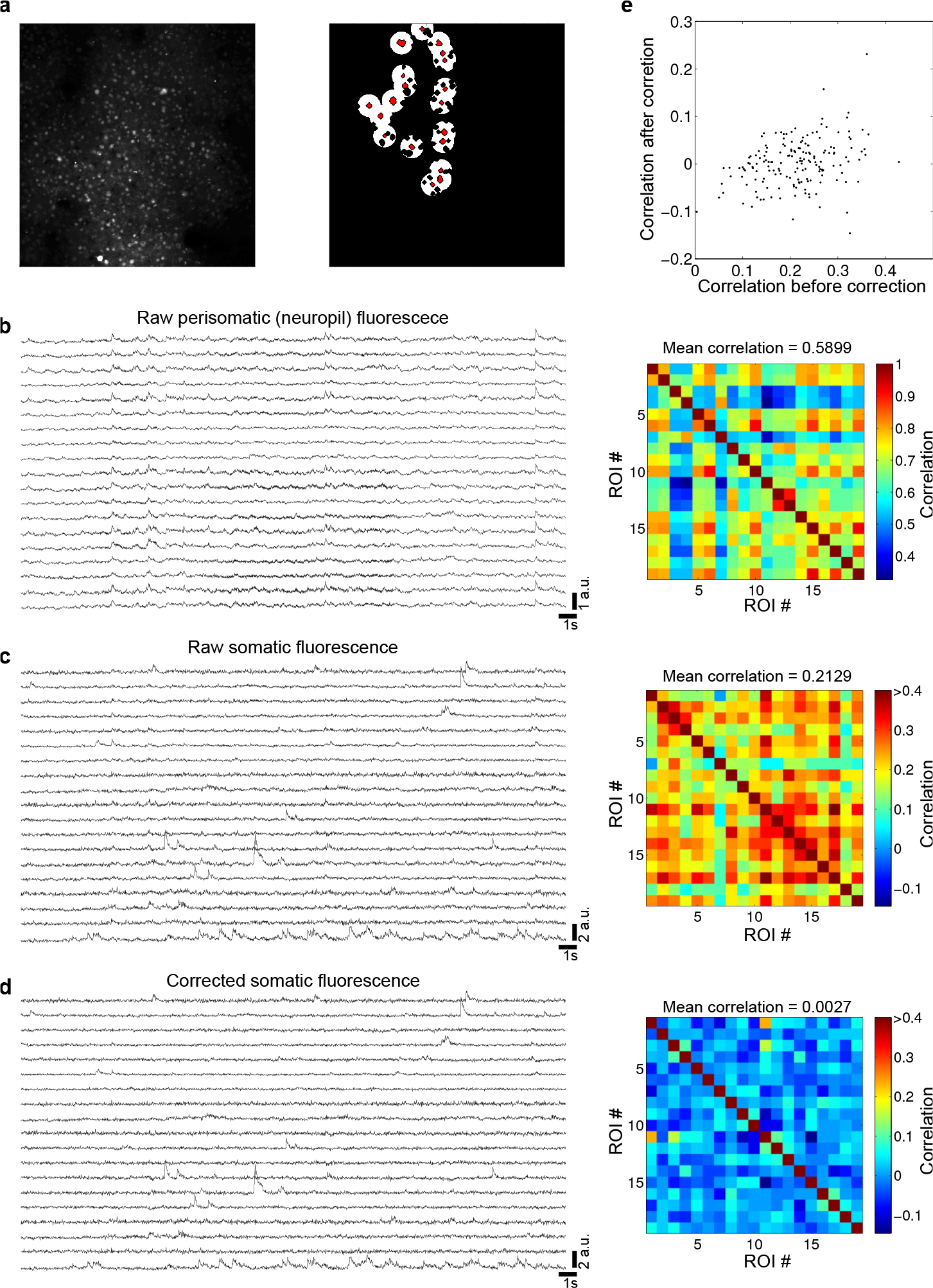
Correction of neuropil fluorescence contamination. **(a)** Left: optical plane imaged with two-photon microscopy of the mouse somatosensory cortex (same data as Fig. 2b). Right: examples of the detected ROIs (red) and their circular perisomatic masks used to calculate the local neuropil signals (white). Perisomatic masks are overlapping, but each ROI is associated with a single circular mask. Black holes inside the perisomatic masks are other detected ROIs not included in the local neuropil signal calculation. (**b**) Left: raw fluorescence traces obtained with the perisomatic masks shown in **a** (neuropil signal). Right: pair-wise correlation matrix for the signals shown in the left. Note the high temporal correlation across the traces. (**c**) Same as **b**, for the raw fluorescence traces of the ROIs shown in **a** (somata). (**d**) Same as **c**, for the corrected ROI fluorescence traces, obtained by subtracting the traces shown in **b** from the corresponding traces shown in **c**, with **α**=0.9. Note the reduction in the temporal correlations, compared to those found in **c**, despite the small changes of the individual fluorescence traces. (**e**) Relationship between the pair-wise correlations shown in **c** and **d.**

**Figure 4.**
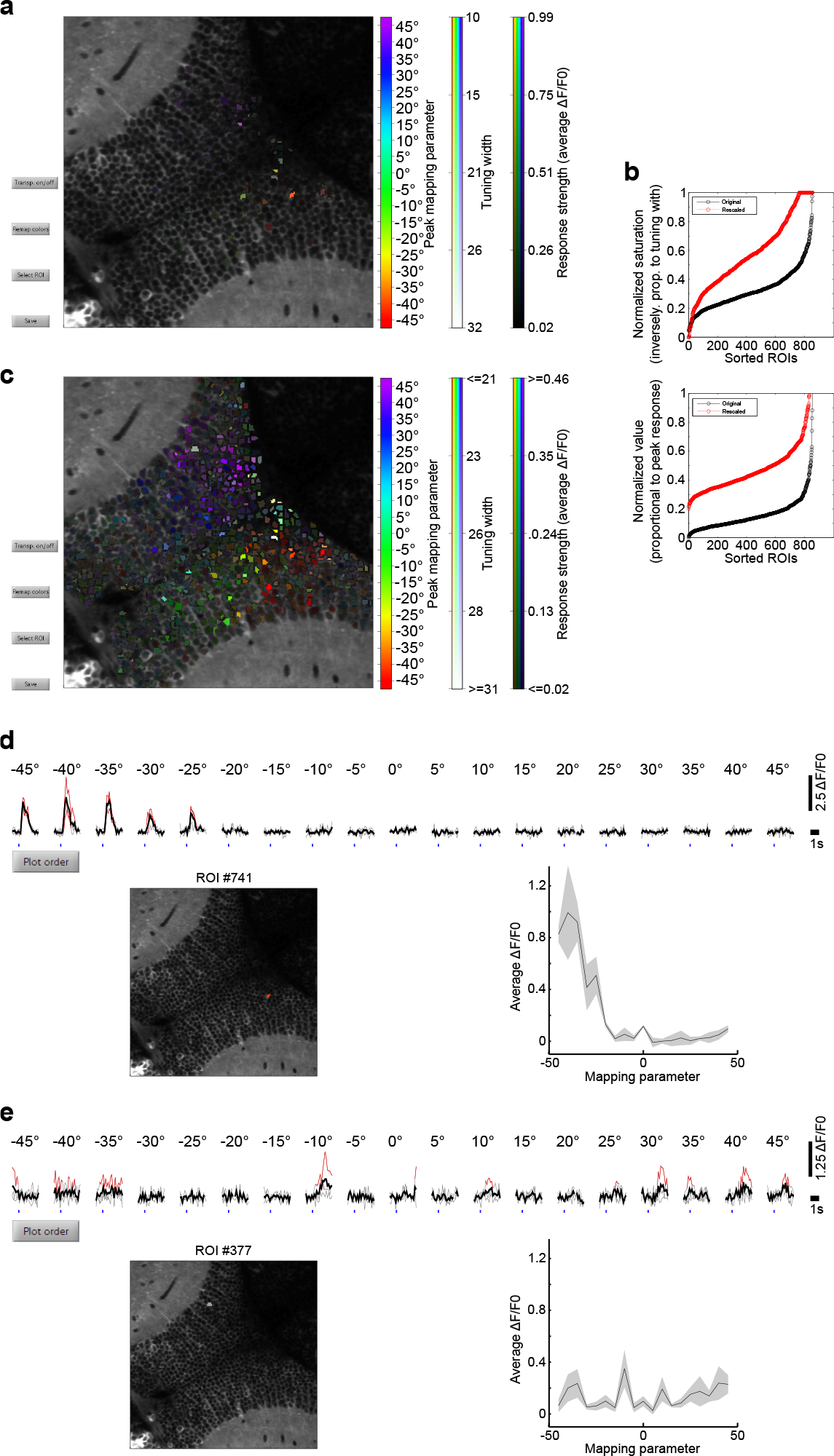
User interface Screenshots of the calculated neuronal responses to experimental events. The larva was visually stimulated with light spots at different visual field azimuth angles. (**a**) ROIs are colored with an *HSV* color code representing their preferred azimuth angle *(Peak mapping parameter; hue)*, azimuth selectivity *(Tuning width; saturation)* and average response at preferred azimuth *(Response strength; value)*. The full *HSV* channel range was used. Due to the skewed distribution of the responses, only a few responsive ROIs can be visualized. (**b**) Offsetting and clipping of the *saturation* and *value* channels to improve visualization. Black, original values used in **a;** red, rescaled values used in **c.** (**c**) Same data shown in **a**, but with the rescaled channel ranges. After this step, the retinotopic organization of the OT becomes evident. (**d** and **e**) Screenshots of the responses of two ROIs selected by clicking on *Select ROI* in **c.** Average (black) and single-trial (gray) *∆F/F0* responses are organized according to the stimulus values, where significant trial responses are shown in red. Bottom right, tuning curves of the ROIs. Black, mean response; gray patch, standard error. Note how the responsive but less selective ROI in **e** is shown with a more whitish color code (low *saturation*).

**Figure 5.**
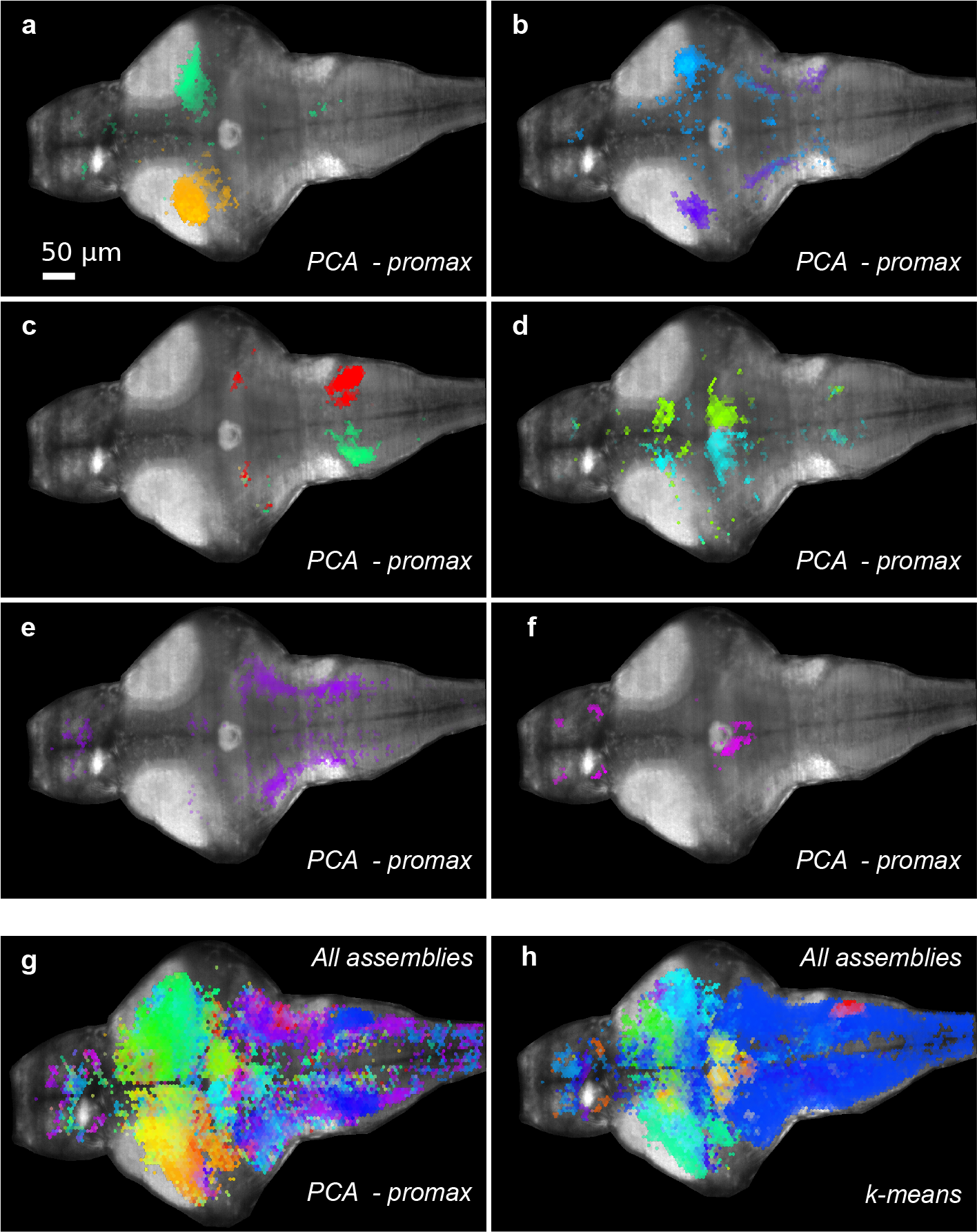
Detection of assemblies of correlated ROIs in zebrafish whole-brain light-sheet imaging reveals the spatial structure of brain activity. Example of a transgenic zebrafish larva pan-neu-ronally expressing GCaMP5 *(HuC:GcaMP5)*. The larva was vol^-^umetrically scanned with fast multi-plane single-photon light-sheet microscopy. 40 optical sections, separated by 5 μm steps in the axial direction, were imaged almost simultaneously at 2.1 Hz. Due to the lack of single-neuron resolution in this imaging dataset, a grid of hexagons (9 Mm in diameter representing approximately the area of an average neuron) was imposed over each imaged optical plane to obtain a total of >40,000 ROIs laid out over the entire nervous system. (**a-f**) Some of the ROI assemblies found with the *PCA-promax* approach, displayed over the maximal intensity projection of all imaged optical sections (for individual optical sections see **Supplementary Video 1**). (**a-d**) Pairs of symmetric unilateral assemblies. (**e-f**) Single bilaterally symmetric assemblies. (**g**) All ROI assemblies found with the *PCA-promax* method, displayed over the maximal intensity projection of all imaged optical sections (for individual optical sections see **Supplementary Video 2**). In all panels, assemblies are displayed with the same color code, according to the similarity of their activity dynamics (the more temporally correlated, the more similar the color; except for the temporally correlated assembly pair shown in **c**, which was colored differently to facilitate visualization). (**h**) Same as **g**, but for assemblies found using *k-means*, color-coded according to the similarity of their activity dynamics. To allow for a comparison between **g** and **h**, the dimensionality of the dataset was reduced through PCA before clustering (otherwise, clustering did not converge). A correlation metric and a *complete* linkage were used. Note how this clustering reveals assemblies whose spatial organization is roughly consistent with those shown in **g**, but exposes a much less biologically relevant fine structure.

**Figure 6.**
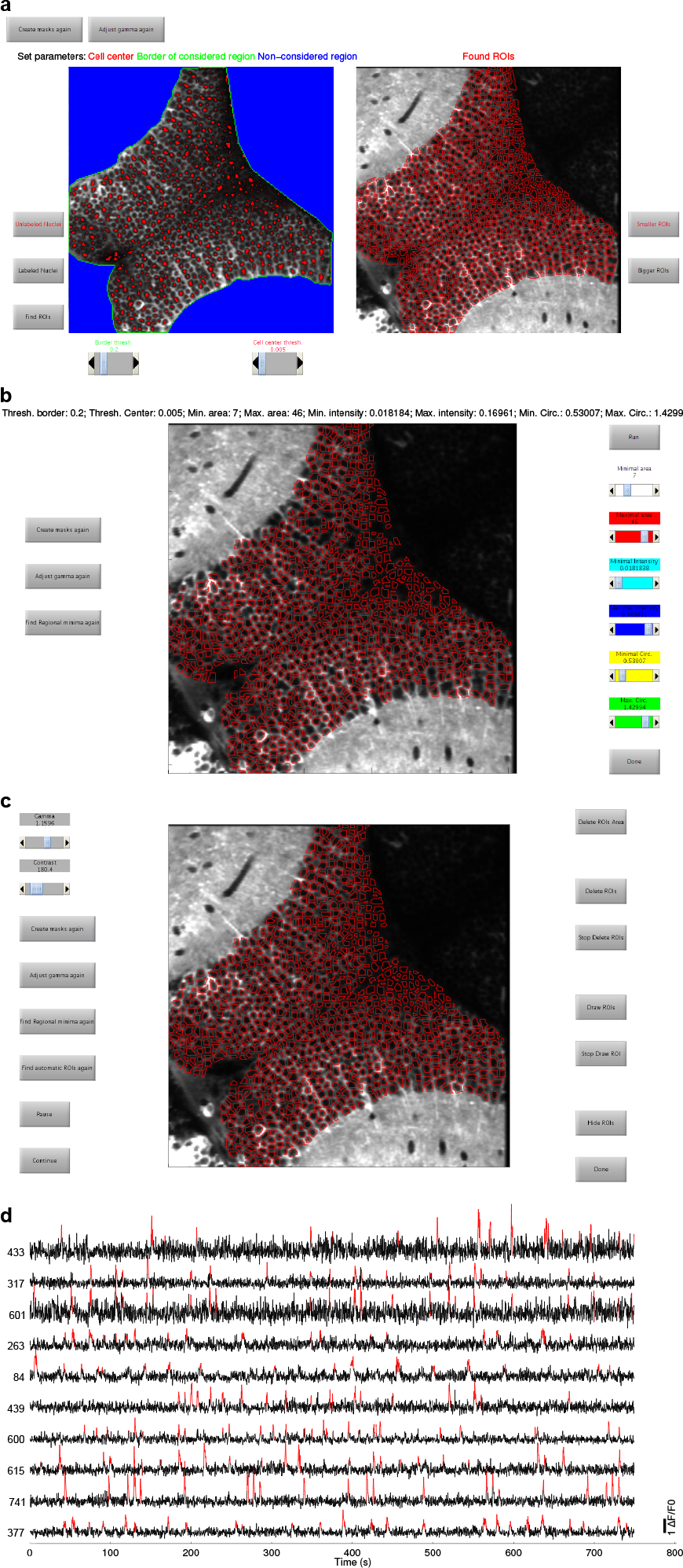
User interface screenshots of three different steps of the ROI segmentation and *∆F/F0* calculation procedure. Optical sections correspond to the OT of a zebrafish larva pan-neuronally expressing GCaMP3 *(HuC:GCaMP3)*. Parameters displayed are those chosen for the case study. (**a**) Left: selection of the *thr_soma_* and *thr_Neuropil_* threshold values for detecting, respectively, putative cellular nuclei (red) and marking regions to be ignored by the segmentation procedure (blue; interface is shown in green). Since neuronal somata are mostly packed in the OT’s *stratum periventriculare* (SPV) layer, the tectal neuropil and surrounding regions were ignored by the mask drawn in step 12. When somata are sparsely scattered in neuropil (e.g., Figs. 2a,b) the latter can be marked with a higher *thr_Neuropil_* value. Right: Perimeters of the automatically detected ROIs (red). (**b**) Selection of morphological criteria to filter automatically detected ROIs (step 18). Red, perimeters of selected ROIs after filtering. (**c**) Final manually curated ROIs (step 18) of the SPV layer. (**d**) Screenshot of the automatically displayed examples of *∆F/F0* traces (black) and their significant fluorescent transients detected (red; step 30).

**Figure 7.**
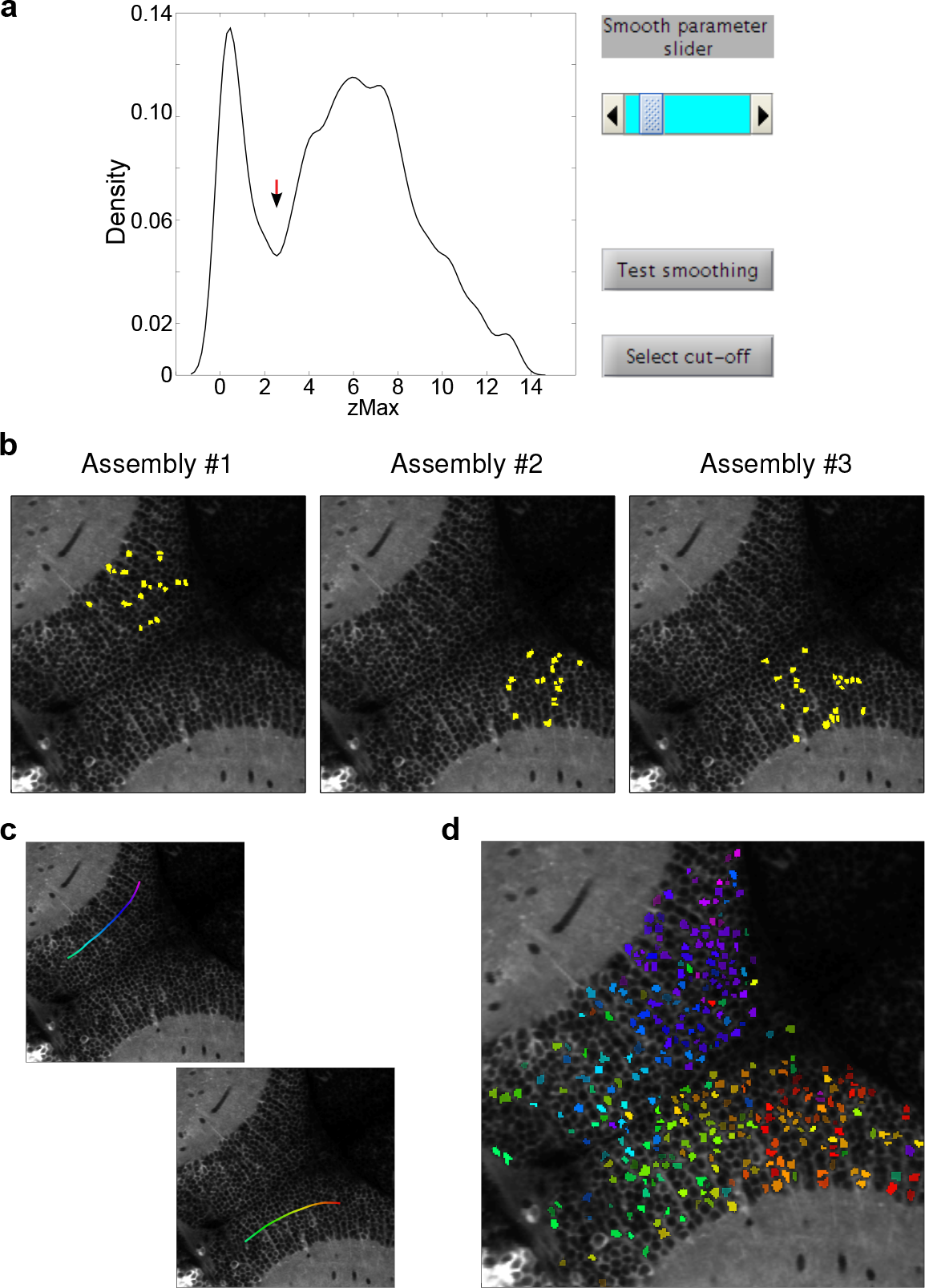
Detection of neuronal assemblies. (**a**) Screenshot for the selection of the *zMax* threshold. After setting the smooth parameter with the slider (top-right), the threshold is chosen with a mouse click on the graph of the density distribution (red arrow). (**b**) Screenshot of 3 of the assemblies (out of 42) automatically displayed. ROIs that belong to each assembly are labeled in yellow. (**c**) Screenshot of the two anatomical axes defined by the user (step 50). The *Along curve* option was selected, two masks were drawn to divide the imaged plane in two regions, and the user drew a curve with the mouse for each region. Each curve is automatically colored such that the combined curves reproduce the hue gradient used in Figs. 4a,c. The chosen curves span the rostro-caudal retinotopic axis of each OT hemisphere. (**d**) Screenshot of the figure obtained displaying the spatial organization of the assemblies along the selected axes. Assemblies’ ROIs are colored according to the defined axis (i.e., the position of the assemblies’ spatial cen-troid with respect to the defined axis). The comparison with Fig. 4c confirms that assemblies reproduce the OT’s retinotopic functional map.

**Figure 8.**
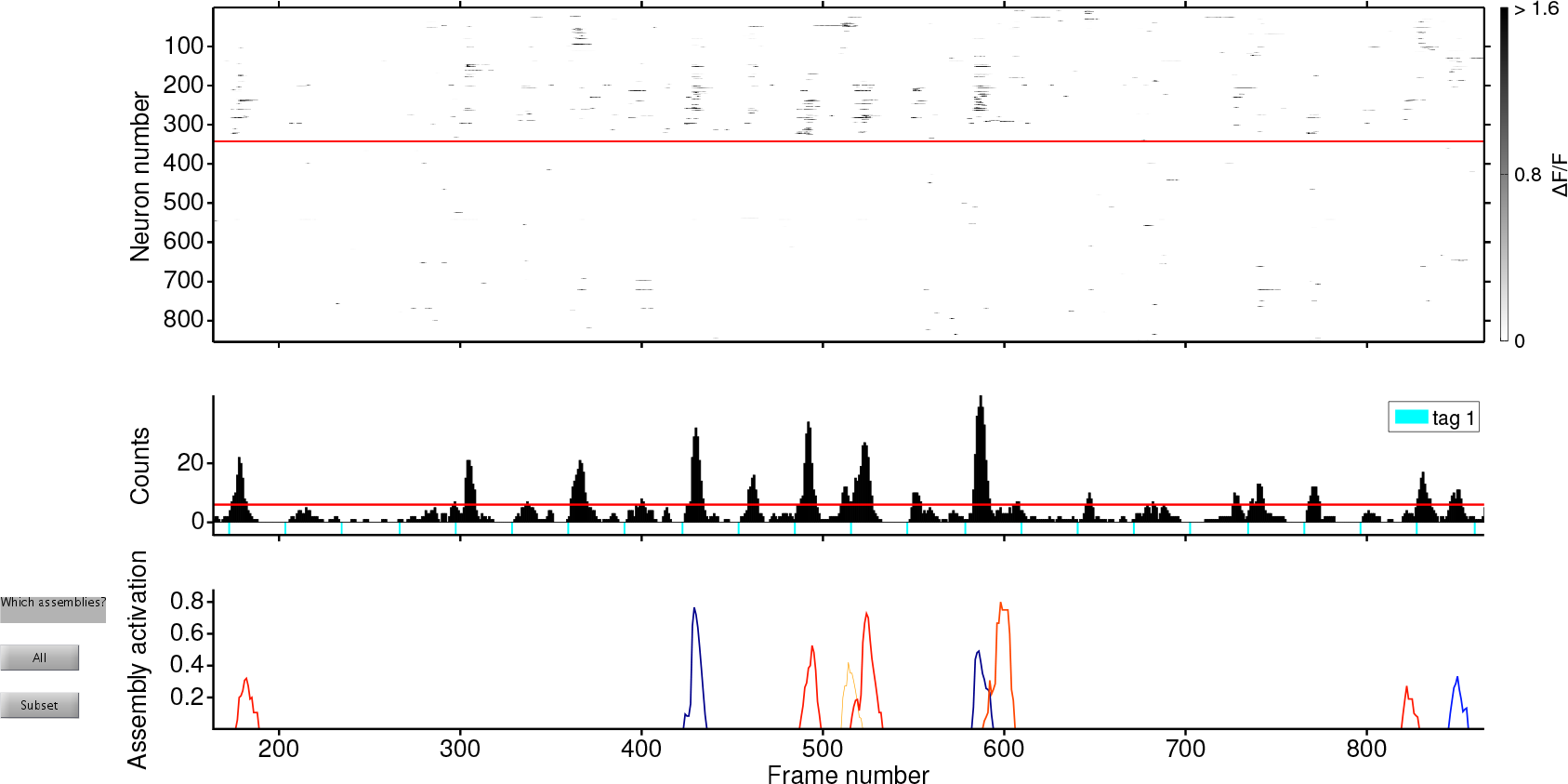
Screenshot of the exploration of the assemblies’ activity dynamics (step 52), zoomed over a 175 s period of the imaged activity. Top: raster plot of the ROI’s significant fluorescence transients, with ROIs sorted according to the order stored in the *_ORDER_TOPO.mat* file, allowing the visualization of distinct assemblies. ROIs not assigned to any assembly lie below the red line. Grayscale: *∆F/F0* amplitude. Middle: Histogram of significant transients in the complete population. The red line marks the threshold for significant population events (peaks exceeding this threshold cannot be explained by chance, as inferred by a shuffling procedure). Color-coded bars below the histogram show the experimental time tags informed in step 51 (tag 1; here, when the larva was visually stimulated). Bottom: Assembly activation dynamics (as measured by the *Matching index*) for 5 assemblies selected with the *Subset* button (same color labels as in Fig. 7d). Note that each activation peak of the same color corresponds to the activation of roughly the same group of neurons.

**Figure 9.**
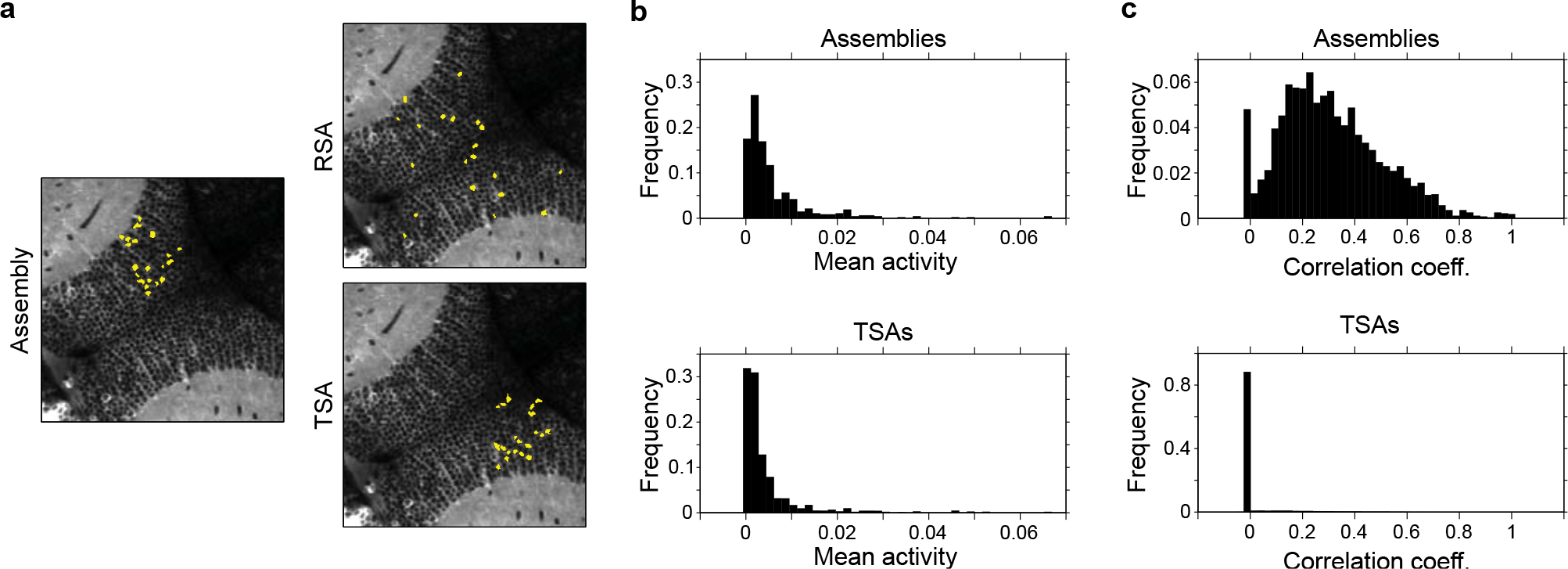
Comparison of specific features of the assemblies with those of surrogate controls. (**a**) Spatial layouts of a given assembly detected in the data, and of one example of a *Random surrogate assembly* (RSA, where an equal number of ROIs are randomly placed) and a *Topographical surrogate assembly* (TSA, where ROIs are placed preserving the inter-ROI distances of the original assembly). (**b**) The normalized frequency histogram of average ROI activity levels (mean significant *∆F/F0* per imaging frame) obtained for ROIs included in assemblies (top) and those included in *TSAs*. (**c**) Same as **b**, but for the ROI activity correlations (pair-wise Pearson correlation coefficient).

For all modules, we provide the source code implemented in *MATLAB* functions and scripts, which the user can adapt and modify, if needed. Given the large scope of the pipeline, in this section we limit discussion to an initial conceptual overview of each module, but we encourage users to gain a better appre ciation of the toolbox through hands-on protocol experience with the provided case study (see *Results*). Most of the underlying mathematical details have been extensively described in Romano *et al*^9^.

### The pre-processing module

This module has several sub-modules that we describe in this section.

#### Drift correction

Longitudinal imaging experiments are prone to present in-plane *x-y* drifts of the imaged optical section. This drift has to be compensated for to obtain stable imaging recordings. This compensation is carried out by the first sub-module (step 1). The series of image stacks is processed with the effective and widely used *Template Matching* image registration plugin of *ImageJ*. Nevertheless, we often observed that this plugin could fail at registering imaging videos that show episodes of large fluorescence variation associated with global neuronal activations. For situations like this, we included an additional program that solves this issue (see *Troubleshooting* in step 1).

#### Artifact detection

*In vivo* imaging may present image frames with motion artifacts produced by diverse sources, like animal movements, respiratory or cardiac/vascular pulsation, etc. The following sub-module (steps 2-7) detects and discards these aberrant frames. First, a program automatically detects a series of candidate frames with movement artifacts by performing a cross-correlation across the imaging frames and an image template. The template consists of a spatially smoothed image frame that preserves the major anatomical landmarks, with reduced image noise (this is particularly important for laser-scanning imaging techniques, like two-photon microscopy). The second part of this sub-module (steps 8-9) is an interactive graphical user interface (GUI) that allows for manual inspection and curation of each artifact candidate. However, if complete automatization is desired, the GUI has the option to skip the individual inspection of each candidate.

#### ROI segmentation

The following sub-module uses digital image processing techniques to perform morphology-based image segmentation, in order to separate the ROIs that correspond to individual cells from the signal of other cells and the surrounding neuropil (steps 10-19). As expected, the success of this module depends on the image’s spatial resolution, the anatomy of the imaged region, and the cellular labeling by the fluorescent reporter. Here, we demonstrate that it can successfully process remarkably different datasets (see below and Figs. 2 and 6). Nevertheless, if the user is not satisfied with the results obtained, the automatic ROI segmentation can be bypassed, since the module’s GUI can also be used to manually draw all the ROIs, to automatically implement a custom hexagonal grid of ROIs (see Supplementary Fig. 1a and Fig. 5), or import predefined ROIs.

The automatic detection of ROIs is performed in a few simple and interactive steps in the GUI, which are explained in the next paragraph. First, the image of the optical plane is spatially normalized. Then, two thresholds, *thr_soma_* and *thr_neuropil_* must be set to obtain the automatically detected ROIs. The GUI then allows the user to curate the resultant ROIs. For this, a series of ROI morphological criteria (area, fluorescent intensity, and perimeter circularity) can be applied to rapidly filter undesired ROIs (step 18; see Fig. 6b). Finally, the GUI allows the user to manually add, delete or modify the ROIs (step 19; see Fig. 6c). We now explain the calculations involved in these steps. Synthetic calcium dyes label cells rather uniformly. On the other hand, genetically encoded calcium reporters are typically expressed in the cytosol, and can be excluded from nuclei or not. Imaging cells where the reporter is excluded from the nuclei results in characteristic ring-shaped fluorescent labels; where the reporter is present in nuclei, imaging results in uniformly filled spot labels. Therefore, this sub-module allows the user to specify the kind of fluorescent labeling used (i.e., labeled nuclei or not), switching between algorithms tailored to detect ROIs in either of these two conditions.

The steps of the ROI segmentation algorithm are the following. First, the imaged video is averaged across frames to obtain the image *avgImg*. Fluorescent labeling may be uneven in the imaged plane. Therefore, the intensity of *avgImg* must be spatially normalized to avoid biases in the segmentation. For this, we first filter *avgImg* with either a 10% or a 90% order-statistic filter (with a running square window), obtaining *avgImg_10_* and *avgImg*_*90*_, respectively. We then calculate the spatially normalized image *avgImg_Norm_* as *avgImg_Norm_* = (*avgImg – avgImg_10_*) / (*avgImg_90_* – *avgImg_10_*). The user can set the size of the square running window so that *avgImg_Norm_* clearly shows the fluorescent-labeled cells with an intensity that does not vary significantly across the imaged plane. If cell nuclei are unlabeled, *avgImg*_*Norm*_ is converted to its complement (i.e., its negative), converting the unlabeled nuclei into bright spots. The user then has to interactively set two threshold parameters, *thr_soma_* and *thr*_*Neuropil*_, that will result in binary masks that identify cell somata and the regions that surround them (i.e., the neuropil), *Mask_soma_* and *Mask_Neuropil_*, respectively (see left panel in Fig. 6a). *Mask_Neuropil_* is calculated through a simple thresholding procedure on *avgImg*_*Norm*_ using *thr*_*Neuropil*_*. Mask*_*Soma*_ is calculated through an extended-maxima transform on *avgImg*_*Norm*_, which consists of the regional maxima of the *H*-maxima transform of *avgImg_Norm_* (obtained by suppressing all maxima in *avgImg_Norm_* whose height is less than *thr_soma_*). Then, both *Mask_soma_* and *Mask_Neuropil_* are imposed as regional minima to the image complement of *avgImg_Norm_*. Finally, a watershed transformation is performed to obtain the ROI perimeters. The imposition of regional minima in *Mask_Neuropil_* prevents the watershed algorithm from finding “catchment basins” (i.e., ROIs) in unwanted regions (i.e., the neuropil). This allows the implementation of the watershed transform in imaged regions where labeled cells are spatially distributed in a sparse or scattered manner. Finally, the GUI can be used to manually curate ROIs, as previously explained.

We provide 5 examples of single-neuron ROI detection performance obtained with two-photon imaging: *i*) for an injected synthetic dye that labels the entire volume of the neurons (OGB-1 AM) in mouse visual cortex^21^ (see Fig. 2a); *ii*) for mCherry-labeled nuclei in a GCaMP6s imaging experiment obtained by viral injections in the mouse somatosensory cortex^11,22^ (see Fig. 2b); *iii*) for large-field imaging of nuclei-excluded GCaMP3 in transgenic zebrafish larvae (see Fig. 2c); iv) for a preparation similar to *iii* but imaging a smaller field (the provided case study; see Fig. 6); and v) an example of single-neuron ROI detection in single-photon high-acquisition rate (100 Hz) light-sheet imaging of a transgenic zebrafish expressing GCaMP5 (see Fig. 2d). These examples illustrate the algorithm’s performance in settings that differ markedly in neuronal labeling density (examples *iv* and *i* being the most and less dense, respectively), and with imaging techniques of different spatial resolution and acquisition rates (two-photon laser scanning and single-photon light-sheet microscopies). As explained in *Results*, we typically obtain a >85-90% success rate in automatically detecting single-neuron ROIs.

#### Calculation of relative fluorescence variation

The final sub-module (steps 20-30) calculates the ROIs’ relative fluorescence variation *(∆F/F0)* and automatically detects significant fluorescence transients in a completely automated manner, based on a few experimental parameters that the user must set. It involves performing an (optional) signal correction from neuropil fluorescence contamination, a data sanity test, and the detection of the baseline ROI fluorescence.

Microscopy techniques are continuously improving in resolution power. However, even two-photon fluorescence microscopy has relatively limited resolution (especially axially) for single-neuron *in vivo* imaging. Thus, the neuropil fluorescence signal can substantially contaminate the somatic signal^23,24^ so that *F_measured_* = *F_soma_* + α *F_neuropil_*. To correct for this, the module allows the user to set the parameter α so as to subtract a local perisomatic neuropil signal from the measured signal^11^ (step 23). To that end, for each ROI, the corresponding local neuropil signal is automatically obtained from a circular area with a 20 μm radius that surrounds the ROI in question and excludes all other ROIs (Fig. 3a). Neuropil subtraction is particularly important when imaging loosely scattered neurons surrounded by neuropil (e.g., Fig. 2a,b). Indeed, signal contamination from the neuropil can notably affect the measurement of the correlations between the neuronal fluorescence time courses (Fig. 3b-e). However, the appropriate value of α is still a mater of debate^25^.

The module then performs a data sanity test that discards ROIs whose fluorescence signal is too low and/ or presents artifactual fluorescence traces (step 22). For the detection of these artifacts, the module first calculates for each ROI a smooth estimate of its slowly varying fluorescence baseline *(F_smooth_)*. This baseline reflects slow fluctuations unrelated to the faster calcium transients associated with neuronal activations (Supplementary Figure 1b,c), and it is calculated using a running-window average of the 8th percentile of the ROI’s fluorescence^26^, with a time window 40 times larger than the decay time constant of the calcium reporter (τ). This procedure results in a *F*_*smooth*_ that robustly tracks the ROI’s basal fluorescence level, without being affected by the fast activation transients. Thus, if an ROI is associated with a *F_smooth_* that shows sudden variations exceeding a user-selected threshold, the ROI is discarded (in practice, this eliminates ROIs of unhealthy neurons or healthy neurons that move in or out of focus during the imaging session). For the calculation of the ROIs’ *ΔF/F0*, the user can choose to estimate *F0* in two ways (step 24): *i*) use the average ROI fluorescence in a user-selected time window (typically, a time window immediately before a particular experimental event, e.g. sensory stimulation, animal movement, etc.); *ii*) use *F_smooth_*. Option *i* is useful for short imaging sessions with clearly defined experimental events, while *ii* is particularly useful for rather long sessions where *F_0_* can vary over time.

#### Inference of significant fluorescence transients

This sub-module automatically infers which fluorescence transients are significantly associated with neuronal activations. A few options are available, and users can choose the inference procedure that best suits their imaging experiments. This is a critical step for the subsequent analysis. Therefore, while this sub-module implements an algorithm for the detection of significant transients, we again allow users to bypass this algorithm and import results from alternative methods^27–29^, if preferred (step 29).

To infer the statistical significance of fluorescence transients, the algorithm implemented in the toolbox considers that any event in the fluorescence time series data belongs to either a neuronal activity process or to an underlying noisy baseline. Thus, the sub-module first performs a key step for this inference, which is the estimation of the ROIs’ fluorescence noise levels (i.e. the baseline fluorescence noise scale σ). The sub-module allows for the estimation of σ using two options (step 25): 1) by fitting a Gaussian process to the negative *∆F/F0* fluctuations of each ROI (i.e., those below baseline, which are not related to calcium transients and hence are due to measurement noise; Supplementary Figure 1d) and estimating the standard deviations of the baseline Gaussian process; 2) by estimating the standard deviations of the ROI’s *∆F/F0* traces after excluding the largest *∆F/F0* excursions (which should represent the calcium-transient peaks). Finally, we provide two options for performing the inference of significant transients (steps 26). First, the user can choose a straightforward method that uses a static fluorescence threshold for each ROI. Only those fluorescence excursions that exceed a multiple of the ROI’s are considered significant. Second, alternatively, the user can adopt a more complex method (but more robust against false positives), which exploits the estimated model of the underlying noise and applies a dynamic threshold that depends on both the ROI’s σ and the biophysics of the calcium reporter. For this, the module implements a Bayesian odds ratio framework that analyzes fluorescence transitions across imaging frames. It labels as significant those transitions whose dynamics meet two conditions: *i)* they cannot be explained by the underlying fluorescence noise, according to a user-selected confidence threshold; *ii)* they are compatible with the reporter’s τ. Finally, the module automatically displays the *∆F/F0* traces and the significant transients found (step 30; Figs. 2 and 6d).

### The module for analysis of responses

Several studies aim to relate neuronal activity to experimental events (e.g., sensory stimulation, behavioral response, etc.). Indeed, imaging is particularly suited for revealing the spatial distribution of neuronal responses (i.e., topographic sensory/behavioral functional maps), which may be crucial to understand brain coding strategies. This module (steps 31-37) associates ROI fluorescence responses with a given experimental variable, and visually displays them on the imaged optical plane to assess their topographical organization. This module can also be used in a stand-alone manner, independent of the previous modules of the processing pipeline, since the user can import *∆F/F0* and ROIs obtained using other procedures (see Fig. 1; step 31). For the topographic mapping of ROI responses, there must be a continuous parametric relation between the user-provided variable being mapped and the experimental event (e.g., the frequency of an auditory stimulus when mapping tonotopicity, the position of a visual stimulus when mapping retinotopicity, etc.).

The user first needs to provide timing information corresponding to the events of interest (e.g., the time of the stimulus), and the module automatically isolates, regroups and displays event-locked single-trial and trial-averaged ROI *∆F/F0* responses (see Figs. 4d,e). If significant fluorescence transients were inferred in the previous module (or through other methods and imported), they will be highlighted, allowing for the evaluation of ROI activations at the single-trial level. This automatic ordering and regrouping of trial responses is particularly useful for mapping stimuli-induced responses, since studies usually randomize trial ordering to avoid neuronal adaptation to the stimuli. Moreover, the information in these trial responses is also summarized in corresponding ROI “tuning curves”, by plotting the trial-averaged ROI *∆F/F0* responses during a time window locked to event onset as a function of the variable (see bottom right panels in Figs. 4d,e).

Finally, these ROI tuning curves are color-coded and superimposed on the imaged optical plane (see Fig. 4c). For this, the algorithm uses a *hue-saturation-value (HSV)* colormap (see Supplementary Fig. 2a), where *hue* represents the variable value *v_peak_* that corresponds to the tuning curve peak (the ROI “preferred” variable value), *saturation* depicts the tuning width around *v_peak_* (the ROI specificity for *v_peak_*), and *value* represents the actual *∆F/F0* value at *v*_*peak*_(*∆F/F0*_*peak*_; the ROI response strength). If the calculated ROI responses are associated with a single type of experimental event (e.g., repeated auditory stimulation with a single tone frequency), *saturation* is not used in the color code, since response specificity cannot be defined. As mentioned above, for this *HSV* color code to make intuitive sense, the variable must parametrically relate to the experimental event. In some cases, the distribution of ROI response intensities may be highly skewed (a few very strong responses dominating a set of weaker ones). This can hinder the appropriate visualization of the data, if the full range of responses is represented without any color rescaling procedure (e.g., Fig. 4a). Therefore, the module allows clipping (to flatten signals that exceed a threshold) and offset of the *saturation* and *value* channels, improving data interpretability (see Fig. 4b; compare Fig. 4c with Supplementary Fig. 2b, where the *saturation* and *value* channels were not taken into account). Moreover, the user can choose to inversely relate *∆F/F0_peak_* to a transparency channel, which smooths out the weak and noisier responses, highlighting the most significant ones.

### The module for detection of assemblies

As for the response-analysis module, this module (steps 38-47) can also be used as a stand-alone module independent of the previous processing pipeline (see Fig. 1). To define the assemblies (clusters of ROIs with similar activity dynamics), the module can detect correlated ROIs by taking into account the unfiltered *∆F/F0* traces or by focusing only on the significant *∆F/F0* transients. Furthermore, it can be used to analyze single-plane (see Fig. 7) and multi-plane volumetric imaging experiments (see Fig. 5). The module has two sub-modules. The first and most important (steps 38-46) implements data clustering. This sub-module applies three different clustering algorithms: *i) PCA-promax clustering; ii) k-means clustering; iii) hierarchical clustering*. Both *ii* and *iii* are standard and widely used clustering methods^30,31^, and *i* was introduced and described in detail in Romano *et al*. 2015. Here we briefly describe these, emphasizing the more novel *PCA-promax* approach.

*PCA-promax* implements a fully automated method that searches for ROIs whose activations are correlated on average along the entire experiment, but it is also capable of defining significant ROI clusters episodically activated in synchrony. Since a given ROI could belong to several functional groups, the algorithm is tailored to define non-exclusive ROI assemblies (i.e., it allows for potential overlap between the detected clusters).

The procedure relies on previously proposed techniques^18,32^. Briefly, it consists of two processing steps. First, it z-scores the activity of each ROI and reduces the dimensionality of the complete z-scored dataset of ROI activities through principal component analysis (PCA). The initial z-scoring homogenizes the variance across ROIs, allowing PCA to reveal the global structure of ROI activity covariance. To define the assemblies, it then uses a second algorithm to partition this space of reduced dimensionality, by means of non-orthogonal factor rotation, *promax^33^*. This latter step extends the simpler PCA-clustering method^32^ to non-exclusive assemblies.

Dataset dimensionality reduction is obtained by only keeping principal components (PCs) with eigenvalues greater than λ_*max*_, a theoretical lower bound to the eigenvalues of informative PCs given by the Marčenko-Pastur distribution^32,34^ (equation 1)

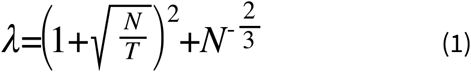

where *N* and *T* are the number of ROIs and imaging frames, respectively. In practice, for long-term imaging data (30 min videos sampled at ∼5 Hz), this procedure typically reduces data dimensionality by ∼100 fold.

As mentioned previously, since we are looking for non-exclusive assemblies, we relax the PCA orthogonality condition^18^. Therefore, to delineate the assemblies, the algorithm works on a space of obliquely rotated components *(promax)*, that sparsely concentrates the PC loadings along non-orthogonal rotated PCs. Hence, after standardizing the loadings on rotated PCs by means of a z-score, a given ROI is included in a particular assembly defined by a rotated PC if its z-scored loading on that rotated PC exceeds a threshold value. This threshold is easily estimated as the first clear minima in the distribution of z-scored maximal ROI loadings (defined by the user at step 42; see Fig. 7a). The algorithm then merges clusters determined by two rotated unitary PCs if the dot product of the latter exceeds 0.6 (i.e., clusters with highly similar neuronal compositions; in practice a 0.6 value identified assembly pairs with exceptional overlap). As a final constraint, only significantly correlated and synchronous clusters are kept (*p* < 0.05, compared to surrogate control datasets (see *Random surrogate assemblies* in Box 1). The assemblies obtained are then automatically displayed.

Briefly, *k-means* partitions data in a fixed number of clusters *k*, defined by the user. It involves randomly selecting *k* initial centroids and assigning each point (e.g., each neuron) to their closest centroids, thus forming *k* preliminary clusters. The centroids are then updated according to the points in the clusters, and this process continues until the points stop changing their clusters (i.e., convergence of the centroids). Typically, the clusters defined by *k-means* are highly independent and uncorrelated (i.e., they present low inter-cluster correlations).

On the other hand, (agglomerative) *hierarchical clustering* groups data by creating a cluster tree or dendrogram. The tree represents a multilevel hierarchy, where clusters at one level are joined as clusters at the next level. First, the similarity between every pair of variables (e.g., neuronal fluorescence traces) is calculated. Then, these similarities are used to determine the proximity of variables to each other. Variables are successively paired into binary clusters, and newly formed clusters are grouped into larger clusters in a bottom-up manner, until a hierarchical tree is formed. Finally, clustering is performed by determining where to cut the hierarchical tree, and assigning all the objects below each cut to a single cluster. For both *k-means* and *hierarchical clustering*, the user can choose to cluster either the original z-scored dataset or the dimensionality-reduced dataset obtained through PCA according to Equation 1. In contrast to *PCA-promax*, for both *k-means* and *hierarchical clustering*, a number of important parameters have to be set. The user must choose between using a euclidean or a pair-wise correlation metric to calculate distances between variables. For *hierarchical clustering* only, the user must choose if the distance between two clusters is defined by the shortest or longest distance between two points in each cluster *(single* and *complete* linkage clustering, respectively). Being classic methods, the particularities of clustering with these metrics and linkages has been extensively reviewed^30,31^. Finally, to obtain the final clustering for both methods, the user has to set the total number of clusters to look for in the data (*k* for *k-means*, and the smallest height at which a horizontal tree cut leaves *k* clusters for *hierarchical clustering*). The second sub-module (step 47) calculates the time series of the assemblies’ significant activations. For this, it uses a matching index, *MI*^*9,35,36*^. The *MI* is defined as

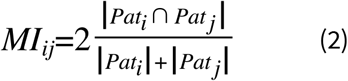

where *Pat_i_* is the binary activity pattern of imaging frame *i*, and *Pat_j_* is the binary target pattern of assembly *j* (i.e., binary *N* × 1 vectors representing the complete population of *N* ROIs, with ones indicating active ROIs and zeros indicating those inactive). Norms are equal to the number of ones in each vector. The *MI* quantifies the proportion of ROI activations that are common to both patterns with respect to the total number of activations present in both patterns. It is valued between 0 (no overlap in activations) and 1 (perfect overlap in activations). To estimate the significance of the assemblies’ *MI*s over the course of the experiment, the algorithm uses the hypergeometric distribution. Under the null hypothesis of independent ROI activations, this is a discrete distribution that describes the probability of having *k* “hits” with *n* target ROI activations in a population of *N* ROIs, showing *K* activations at a given moment. Therefore, it allows estimation of the probability of observing a given activation match by chance, with ROIs independently activated. In step 47 the user can select the threshold *p*-value to consider an assembly activation significant.

### The module for exploratory analysis of assemblies

The first sub-module of this module (steps 48-50) allows for the exploration of the assemblies’ overall spatial (i.e., topographic) arrangement. Individual assemblies can display diverse topographies, and it is informative to analyze whether there is any spatial organization across assemblies. In order to reveal their collective topographic layout, the user can order assemblies according to the similarity of their activity dynamics, or the similarity of their spatial distribution. Irrespective of these two arrangement procedures, assemblies’ ROIs are displayed superimposed on the imaged optical plane, color-coded according to the arrangement obtained.

For the first activity-based arrangement option, the correlation between the assemblies’ *MI*s dynamics are used as a distance measure to build a hierarchical clustering tree of the assemblies’ activity. These distances and the clustering tree are then used to arrange assemblies according to the optimal leaf order that maximizes the similarity between adjacent leaves by flipping tree branches. Colors are then automatically assigned to represent the leaf order. In this way, assemblies with similar dynamics will be similarly colored. This is an unsupervised procedure that does not need user input, and thus it is suitable for automatic analysis of large datasets. An example of this kind of ordering procedure is shown in Fig. 5 for whole-brain light-sheet imaging.

For the second spatial-based analysis procedure, the user indicates an anatomical axis in the imaged plane, and the assemblies will be ordered according to the position of their centroids along the chosen axis. The user can choose the left-to-right or bottom-to-top axis, or draw an arbitrary curve. The user can segment the imaged plane in different regions with different axes (see Fig. 7c). Assemblies that “belong” to a particular imaged region (i.e., the majority of its component ROIs reside in the region) are then color-coded according to their order over the corresponding axis. If there is any particular spatial arrangement along the axis, a clear color gradient should be observed (see *Results* and Fig. 7d).

The second sub-module (steps 51-52) allows an interactive inspection of the neuronal population activity raster plot and the dynamics of assemblies’ activations. The user can import color-coded time tags to highlight the network and assembly dynamics associated with these experimental events (e.g. behavioral epochs, sensory and/or neuronal stimulation times, etc.; see *Results* and Fig. 8).

### The surrogate control module

When validating results obtained from large multidimensional datasets, the use of appropriate control surrogate datasets is critical. This module (steps 53-55) allows pooling user-defined features of neurons within the assemblies (e.g., pooling the average neuronal activation frequencies, the neuronal tuning curves, neuronal phenotypes, pair-wise activity correlations, pair-wise topographical distances, etc.) to assess the statistical significance of the assemblies with respect to surrogate shuffled datasets. Certainly, the choice of control datasets that serve as null models will depend on the particular scientific question being statistically tested. Here, we focus on two particular kinds of control datasets. We preserve the imaged fluorescence dynamics of all ROIs, and regroup ROIs according to: *i)* surrogate shuffled assemblies (where the ROIs of any given original assembly are randomized); *ii)* surrogate assemblies that preserve the original assemblies’ spatial features (see Box 1 and Fig. 9).

## COMPARISON TO OTHER METHODS

### ROI segmentation

ROIs can be determined with algorithms designed to detect either morphological^11,23,37^ or activity^8,28,38^features (e.g., pixels forming round fluorescent spots, or temporally correlated pixels, respectively). These approaches have different advantages and shortcomings. Clearly, the information embedded in the temporal dynamics of fluorescence can be exploited to obtain ROIs. However, activity-based methods are usually prone to target highly active neurons, whose signal-to-noise ratios (SNRs) are strongest. One popular activity-based approach uses independent component analysis (ICA) to parse the multi-dimensional signal into a combination of sparse and statistically independent signals^28^. However, this algorithm becomes impractical for large fields of view with several hundred ROIs, and assuming statistical independence (i.e., decorrelation) of signals can lead to an incorrect segmentation of single-neuron ROIs. Similarly, methods that aggregate correlated pixels^8^ may fuse adjacent temporally correlated neurons (indeed, neighboring neurons in the mouse cortex tend to be more correlated than distant ones^39^). This is particularly problematic for studies of neuronal coding, and clearly limiting when investigating neuronal activity correlations (e.g., when looking for neuronal assemblies).

On the other hand, morphological-based approaches are computationally less intensive and, most importantly, unbiased with respect to the neurons’ activity patterns. The caveat of this approach is that it may result in relatively imprecise ROI perimeters, and could potentially miss ROIs with low baseline fluorescence in weakly labeled imaged regions (for example, due to the spatially non-uniform staining of synthetic dye injections).

Our toolbox implements a more conservative morphological-based algorithm. To avoid segmentation problems of the imaged optical planes with uneven fluorescent labeling, the segmentation is performed on a spatially normalized image *avgImg_Norm_* of the acquired optical plane (see *ROI segmentation* in *The pre-processing module*). Importantly, the morphological algorithms presented here performed very well in the different preparations and the techniques studied (Fig. 2). Nevertheless, the GUI allows the user to skip the automatic ROI detection. In this case, the ROIs can be drawn manually, or a hexagonal grid of ROIs of a desired size can be implemented. Furthermore, the pipeline allows the import of ROIs obtained with other methods, such as the promising spatio-temporal hybrid approaches that are emerging^40^, which seems to perform remarkably well in demixing fluorescent signals from potentially overlapping neuronal sources.

##### BOX 1 Surrogate assemblies for analysis of assemblies’ features

Assemblies consist of groups of neurons (i.e., groups of ROIs). Do neurons that belong to a given assembly share any particular (spatial, molecular, activity-related, etc.) properties? If so, this could indicate an underlying organizing principle. Simply pooling assembly neurons and comparing their features is not enough, especially considering large heterogeneous neuronal populations composed of different sub-populations unevenly represented in the data. One solution for this potential population sampling bias is to statistically test against surrogate control assemblies. For this purpose, for each experiment we created different sets of assemblies shuffled in different ways (see *Random surrogate assemblies* and *Topographical surrogate assemblies* below) and controlled for different properties of the data. The significance of a given assembly feature observed in the data is thus quantified with respect to the surrogate datasets by calculating the probability of obtaining a feature value at least as extreme as the one measured in the data (*p*-value) in the surrogate assemblies.

###### Random surrogate assemblies (RSAs)

For these control datasets, we permute the indexing of the entire neuronal population for each experiment. These permutations allow the creation of a set of shuffled surrogate assemblies for each experiment by using the original dataset clustering (i.e., the original non-permuted neuronal assemblies), but sampling neurons according to the permuted indexes. This procedure randomizes only the spatial topographies of the assemblies, while keeping every other feature of a given dataset intact (e.g., the particular overlaps between assemblies, the number of neurons per assembly, the topographical position and the activation time series of each neuron). Conserving these properties enables testing the specificity of the neuronal associations present in the assemblies.

###### Topographical surrogate assemblies (TSAs)

To test the specificity of neuronal assemblies’ features, while controlling for their spatial layout, we impose on the random surrogate assemblies the additional constraint of conserving the distribution of the relative pair-wise physical distances between neurons in the original assemblies, *p(d)*. To build the TSAs for a given assembly with *n* neurons, we randomly choose a neuron and iteratively add n-1 neurons to the assembly’s TSA in a way that conserves the assembly’s *p(d)* as best as possible. To this end, at a given step *j* of the iteration (when the TSA has already *j*-1 neurons), we add a new neuron whose pair-wise distances to the *j*-1 TSA neurons are most similar (in a mean-squared-error sense) to any given set of *j*-1 distances present in the original assembly. When the iteration of a TSA is complete, we only keep the TSA if its *p(d)* is not statistically different from the original assembly’s *p(d)* (significance threshold is *p*-value>0.05; typically we obtain *p-value*>0.7).

### Inference of neuronal activations from fluorescence transients

Calcium imaging is usually performed to monitor neuronal spiking activity. However, inferring spikes from fluorescence fluctuations is a complicated computational problem, due to the complex relationship between these two variables. The reporter’s fluorescence is a non-linear function of intracellular calcium, and the latter is also non-linearly related to the recent neuronal activity history. However, different fluorescence deconvolution or template matching methods have been proposed to infer estimates of spike trains and spike-rate dynamics^27,29,40,41^. Despite not accounting for the non-linear spikes-fluorescence coupling, these algorithms have been reported to be fairly effective. Nevertheless, their performance critically depends on many experimental factors, requiring parameter tuning that usually involves electrophysiological recordings. Moreover, neurons show diverse relationships between the number of spikes fired and their fluorescence changes^42^, therefore the parameters obtained for one set of neurons will not necessarily apply to other sets of neurons or experiments. Hence, we opted to keep signal filtering as minimal as possible, and implemented algorithms to detect significant fluorescent transients that can confidently be associated with neuronal activity. Importantly, as in deconvolution and template matching methods, our framework allows the extraction of fluorescence transients whose temporal dynamics are compatible with the calcium reporter biophysics (i.e., its decay time constant, τ). Revealing these significant transients is particularly important for detecting neuronal activations in behaving animals on a single-trial basis, where trial-averaging is precluded or sub-optimal^43^, especially when working with low SNR signals.

### Detection of functional assemblies

The identification of assemblies in large neuronal datasets is still an open problem in neuroscience. Template-matching studies have been successful in revealing the activation dynamics of a predefined population pattern^44^. These are supervised learning algorithms, since they work on labeled data (i.e., the given pattern of interest). However, to reveal hidden structure from unlabeled data, unsupervised learning algorithms are necessary. Between these approaches, we can highlight pioneering studies seeking to assess the presence of higher-order (i.e., higher than pair-wise) correlations in neuronal activity^45,46^. However, these approaches only work for triplets in sets of a few tens of neurons, and did not identify the neurons that participated in the correlations. More recently, statistical models have been used to study the prevalence of these high-order correlations in larger networks^47–51^. On the other hand, shuffling methods developed to infer the significance of repeated firing sequences were recently tested on electrophysiological recordings of ∼100 neurons^52,53^. However, these algorithms rely on temporally precise coordination across neurons and, to our knowledge, have never been applied to imaging studies. Promisingly, others^16,17,32^ have applied PCA to successfully isolate assemblies of correlated neurons, by clustering neurons according to their loadings in each principal component (PC). Importantly, the use of the Marčenko-Pastur distribution introduced by Peyrache and colleagues^32^ (equation 1) allows the use of analytical and reliable statistics to infer the significantly correlated signal in the data, instead of the surrogate methods implemented in previous studies^45,46,52–54^. This is important, since there is still no clear agreement on which statistical features should be preserved in these control surrogate datasets.

Despite these important advances, because PCA represents the data in a space of orthogonal PCs, using PCs to delineate assemblies is misleading when neurons participate in different functional assemblies^18^. The *PCA-promax* algorithm thus further extends this work, relaxing the orthogonality condition and using the promax-rotated PCs to isolate assemblies. In this way, the algorithm is capable of detecting non-exclusive assemblies that share ROIs. Furthermore, this approach is data-driven, without the need for any parameter fine-tuning. In contrast, two of the most popular unsupervised clustering algorithms used in the field, *k-means clustering* and *hierarchical clustering* (both also implemented in this toolbox), rely on the definition of key parameters (i.e., the total number of clusters and a distance threshold for the cluster tree, respectively). Furthermore, they bind members (e.g., neurons) to exclusive and non-overlapping clusters, which is problematic considering the distributed coding of the brain, as previously discussed.

Nevertheless, both *k-means* and *hierarchical* clustering (and their variants) have previously been successfully used to detect functional neuronal clusters in imaging experiments^7,8,55–58^. Remarkably, as we have previously shown^9^, the *PCA-promax* algorithm consistently and significantly outperforms both of these methods, as quantified by two standard cluster quality indexes: 1) the *Davies-Bouldin* index, which measures the ratio of the within-cluster scatter and the between-cluster separation; 2) the *normalized Hubert’s* Γ index, which quantifies the agreement between the observed pair-wise neuronal correlations and the ideal case, in which pairs in assemblies have identical activities and pairs from different assemblies are uncorrelated. This improvement is most probably due to the fact that while both *k-means* and *hierarchical clustering* group variables according only to pair-wise distance measures (e.g., pair-wise correlation coefficients, euclidean distances, etc), PCA is designed to reveal the global underlying correlational structure of the dataset through an eigenvector-based multivariate analysis. Furthermore, this improvement could also be related to the previously mentioned fact that *k-means* and *hierarchical clustering* seek to detect independent or hierarchically organized clusters, respectively (see *The module for detection of assemblies*). While the former clustering feature is clearly not in agreement with the distributed nature of brain processing, the latter hierarchical character may be consistent with the hierarchical organization across brain regions, but is probably not compatible with the functional organization at the neuronal sub-circuit level. Importantly, since the *PCA-promax* method involves a dimensionality reduction procedure, it is effective in processing large datasets, as demonstrated by the detection of biologically relevant assemblies in whole-brain zebrafish light-sheet imaging experiments (∼ 40,000 ROIs, Fig. 5). To our knowledge, few studies^7,8,59^ have attempted to perform clustering in these large-scale imaging datasets, and they have all used *k-means* clustering.

As discussed above, functional clustering of neuronal data is a complex mathematical procedure still under development, and the *PCA-promax* method for detecting assemblies has limitations. The algorithm relies on PCA, which implements linear data transformations. Thus, non-linear neuronal activity correlations (i.e., when plotting the activity of a neuron against another neuron produces a curved cloud of points) could result in spurious assemblies. Most multivariate statistical analysis methods currently applied are also linear, but those that rely on manifold learning algorithms^60,61^ and network theory^54^ could accommodate other non-linear measures of activity similarity. Furthermore, when using PCA, there is no guarantee of a straightforward interpretation of the PCs obtained in terms of biological or experimental variables. In principle, this could be improved with the recently proposed demixed PCA technique^62^, which reduces the dimensionality of the data, taking into account task parameters (e.g., sensory or motor variables). Furthermore, increasingly popular non-negative matrix factorization techniques^63^, which unlike PCA impose a positivity constraint to its components, may be more biologically interpretable in terms of positive neuronal activations. Finally, the *PCA-promax* method is particularly suited to delineate discrete clusters of ROIs that are engaged in brief events of collective activation. Thus, for population activations that present a continuous and long-lasting evolution in time (e.g., propagating spatial waves, oscillations), this algorithm would tend to discretize these dynamic phenomena in distinct patterns. For example, a wave would be separated in a progression of distinct assemblies that represent different instances of the moving wavefront. Therefore, particular care should be taken in the interpretation of the results in these kind of scenarios. Nevertheless, this is not a specific problem of the *PCA-promax* algorithm, since time-evolving patterns are usually problematic for all clustering methods.

### Comparison to other toolboxes

There are few software packages available for a comprehensive analysis of calcium imaging data, despite several recent and important developments^40,64,65^. Here, we briefly describe these available toolboxes and compare them to the one presented here.

These available toolboxes are implemented in *Matlab*^*40,64*^ and *Python*^*40,65*^ programs, and they are all focused on pre-processing analysis of imaging data. They include within-plane motion correction for laser scanning microscopy^65^, image segmentation to obtain neuronal ROIs, extraction of raw fluorescence signals^40,64,65^, and signal denoising and spike deconvolution^40,64^. In particular, the most salient feature of the Tomek *et al*. package is its accelerated cell morphology-based segmentation algorithm, which can detect ∼100 ROIs in 256×256 pixel images in a few milliseconds, opening the possibility for online analysis. On the other hand, Pnevmatikakis *et al*. use a powerful spatio-temporal analysis framework based on constrained non-negative matrix factorization, capable of demixing spatially overlapping neurons, which may prove important for tissue densely packed with somata.

Like the above described packages, this toolbox also performs data pre-processing and, as shown here, can be efficiently applied to different fluorescent reporters, imaging techniques, and imaged regions (Figs. 2 and 6). In addition, the present toolbox also covers the correction for neuropil signal contamination, mapping of functional neuronal responses and the analysis of population activity dynamics. Thus, this is the first protocol for a toolbox that covers the complete workflow from raw data to the automatic display of results. For example, Tomek *et al*. presented a supporting module to export fluorescence data along with stimulation time tags, which the user could in turn use to additionally develop a mapping procedure of functional responses. Here, we present an interactive response analysis module capable of automatically producing publication-quality figures, rich in interpretable functional information (Fig. 4). Furthermore, our package performs automatic cluster analysis for the detection of neuronal assemblies, even in whole-brain data (Pnevmatikakis *et al*. only pre-processed and segmented this kind of large-scale data). Finally, it contains modules for interactive and exploratory analysis of the neuronal population activity in the context of experimental and/or behavioral events, which can be extremely useful when dealing with such complex and multivariate datasets.

With respect to its implementations, the toolbox is designed in a flexible manner that allows importing data from other sources, and using separate modules in an autonomous way, bypassing preceding modules if desired. No previous coding experience is required for its use, since interaction with the toolbox is implemented through diverse graphical user interfaces (GUIs). On the other hand, command-based packages^65^ require more coding expertise, but can be run in batch mode without user interaction to automatically process collections of data. Nevertheless, complete automatization typically comes at the expense of a higher rate of data-processing errors, compared to pipelines that incorporate manual curation.

## PREVIOUS APPLICATIONS

This toolbox has been used by previously reported studies for the analysis of calcium imaging data of transgenic GCaMP-expressing zebrafish larvae. We used the toolbox in order to study the spontaneous neuronal activity patterns of large (∼1000) neuronal populations in the optic tectum (OT), by monitoring GCaMP3 fluorescence through two-photon microscopy (4 Hz scanning rate). Through the *PCA-promax* clustering algorithm, we demonstrated that the ongoing spontaneous activity of this sensory brain region was organized in assemblies of functionally similar neurons, which reflected neuronal mechanisms that assure robust circuit functioning for the extraction of behaviorally relevant visual information^9^. Thompson and colleagues also used this algorithm for detecting assemblies from GCaMP5G single-plane light-sheet imaging data of ∼500 simultaneously monitored OT ROIs scanned at 10 Hz, revealing tectal assemblies responsive to different sensory modalities^20^. They further demonstrated that these assemblies are composed of a reliably responsive core of neurons together with a more variable group of partner neurons, revealing a mechanism for robust neuronal coding of stimulus features while retaining contextual flexibility. More recently, the toolbox has been also used to demonstrate that sustained rhythmic neuronal activity among a specific group of direction-selective tectal neurons was associated with the illusory perception of visual motion^71^. These examples demonstrate that the analysis framework implemented in this toolbox can effectively shed light on the neuronal interactions that underlie brain computations.

Finally, in this work, we show that the pre-processing module can also be applied to detect single-neuron ROIs and significant fluorescent transients in the cortex of mice injected with OGB-1 AM (sampled at 30 Hz; Fig. 2a) or expressing genetically encoded GCaMP6s (sampled at 7 Hz; Fig. 2b) imaged through two-photon microscopy, and in transgenic zebrafish larvae imaged through large-field two-photon mi croscopy (GCaMP3, sampled at 1 Hz; Fig. 2c) or single-photon light-sheet imaging (GCaMP5, sampled at 100 Hz; Fig. 2d). Moreover, we show that the module for detecting assemblies is remarkably efficient in finding biologically meaningful assemblies in datasets of multi-plane whole-brain light-sheet imaging in zebrafish larvae, consisting of more than 40,000 ROIs scanned at 2.1 Hz (Fig. 5).

## MATERIALS

### Equipment

The pipeline can be run on any desktop computer, but we recommend multicore computers for shorter computing times, as several algorithms allow for parallelization (especially for the pre-processing module). For analysis of light-sheet imaging data, we used a computer cluster of 328 CPUs based on a HTCondor parallelization system. Imaging videos of long experiments can result in file sizes of several gigabytes (GB), and therefore we recommend 64-bit architectures with at least 12 GB of RAM. We also recommend screen resolutions of at least 1280 × 800 pixels for a better experience when visualizing data and using user interfaces.

A *Matlab* installation (MathWorks) is required, including the following toolboxes: Curve Fitting, Image Processing, Statistics and Machine Learning, and Parallel Computing. This list of toolbox requirements refers to the full processing pipeline; specific modules need different subsets of these toolboxes. We verified compatibility specifically for *Matlab* versions R2010a, R2011b, R2014a, and 2015b, but our code should, in principle, be compatible with any later version.

Computer operating systems: any version of Windows, Linux or Mac OS X compatible with the *Matlab* installation.

### Data files

An example case study is included in the downloadable toolbox distribution. It consists of zebrafish larva *in vivo* two-photon calcium imaging data (see *Results*).

### Software setup

*Installation of the toolbox*. Download the analysis toolbox from github.com/zebrain-lab/Toolbox-Romano-et-al. It contains all the *Matlab* source code and a *Readme.pdf* file that explains all the relevant variables used during the pipeline (this file is vital if the user wishes to adapt or further develop the toolbox). It also contains test data for a case study. Double-click on the downloaded toolbox *zip* file to extract its contents. Add the toolbox to *Matlab* path. For this, open *Matlab* and type the command:

> *addpath(genpath(‘Extracted_Folder’))*

where Extracted_Folder is the name of the toolbox unzipped folder.

*Install ImageJ and its Template Matching and Slice Alignment plugin*. Download *ImageJ* from http://image-j.nih.gov/ij/ and the *Template Matching and Slice Alignment* from sites.google.com/site/qingzongtseng/template-matching-ij-plugin and install them following the installation instructions of the corresponding websites.

## PROCEDURE

### Pre-processing module

1) Video registration. If there are no observed movements in the x-y plane of the imaged optical section, continue with the next step. Otherwise, open the *TIFF* stack in ImageJ, click on the *Plugins* menu, then on *Template Matching*, and then on *Align slices in stack*. Select *Normalized correlation coefficient* as *Matching method*, and click on *OK*. Define a landmark region on a reference frame by clicking and dragging to draw a rectangle, and then click on *OK* to launch the registration. If the video was properly registered, save the registered file by selecting *Save* from the *File* menu.

##### CRITICAL

It is of paramount importance to obtain a properly registered video. If there is a significant residual drift after the registration procedure, ROIs will not label a stable region of the imaged plane, compromising the subsequent analysis.

##### TROUBLESHOOTING

If video registration is unsatisfactory, close the video without saving it, re-open it and run the registration procedure again, starting from a different reference frame or selecting a different landmark region. If you observe registration defects at imaging frames that present strong global fluorescence variation events, click on the *Results* window of *ImageJ*, select *Save as…* from the *File* menu, and save the registration results in a *TXT* file with the same name as the imaging video. Then, run *SmoothRegistration.m* in *Matlab*, and check if the output video (a file with a *_reg_smooth.tif* suffix) is properly registered.
2) Find potential imaging artifacts by running the command *findArtifacts.m* in Matlab, and select the registered *TIFF* file.
3) A figure will display an image of the temporal average of the *TIFF* file, and you will be asked to indicate the regions of the optical plane to be analyzed.
4) Click and drag to draw a rectangular mask that excludes inadequate image borders that could have been produced by the video registration performed in step 1.
5) Draw a polygon mask that excludes imaged regions that are of no interest and could introduce artifacts, if there are any. Finish this mask by right-clicking inside the drawn polygon and selecting *Create mask*.
6) Click and drag to draw a rectangle mask that determines the image template over which the cross-correlation will be calculated across the imaging frames.

##### TIMING

Depending on the size of the imaging stack and the speed of the computer, the calculation of the cross-correlation may take 5-15 minutes.
7) A graph will display the cross-correlation values across frames (blue line) and a threshold level (green line, default value −3). The frames with cross-correlation values lower than the threshold potentially contain an imaging artifact. Press the *Enter* key, and you will have the option to change the default value for the threshold, if desired.
8) A GUI will allow you to manually examine each imaging sequence containing an imaging artifact candidate. If you want to automatically label all candidates as validated imaging artifacts without individually inspecting them, click on *Accept All* and continue in the next step. Otherwise, navigate through the sequences in question by pressing on the *Previous frame* and *Next frame* buttons. Frames under examination for putative artifacts are labeled in red on the *Frame number* display. If you determine that the frame contains an imaging artifact, click on the *Yes* button under the question *Does frame X contain an artifact?* (where *X* is the number of the frame being examined), otherwise click on No. You can pause the evaluation at any time by clicking on *Pause*, and resume the inspection by clicking on *Continue*.

##### TIMING

Typically, each putative imaging artifact can be inspected in the GUI in a few seconds. Depending on their number (which depends on the stability of the imaged video and the threshold defined in step 7) this could take a few minutes.
9) A dialog box will ask you if there are frames that you want to add to the artifacts list. If you do not wish to add any frames, just leave the question dialog blank. The final list of detected artifacts will be saved in a file with a *_ARTIFACTS.mat* suffix.

### Segmentation of morphological Regions of interest (ROIs) and extraction of their fluorescence traces

10) Run the script *FindROIs.m and* select the fluorescence imaging *TIFF* file. Set the image pixel size in the *x* and *y* directions (in μm). To use the algorithm for automatic morphological ROI segmentation, select *Automatically detect single-neuron ROIs* and continue along steps 12-19. To implement a grid of adjacent hexagonal ROIs, select *Use hexagonal grid of ROIs* (proceed to steps 11-12, skipping steps 13-19). Otherwise, to import a pre-defined mask of ROIs or define ROIs through manual drawing, select *Import or manually draw all ROIs*, and then respectively select *Manually draw all ROIs* (and continue with steps 13 and 19), or *Import ROIs* and select a *.mat* file containing the desired ROI masks (see Box 2 for file format; then proceed to step 20, skipping steps 11-19). In all cases, at the end of the selected procedure, all the resulting relevant variables will be saved in a *Matlab* file with a *_ALL_CELLS.mat* suffix to the *TIFF* filename.

##### TIMING

As discussed in *Results*, while the automatic segmentation and posterior manual curation can typically be performed in less than 10 min for up to ∼1000 ROIs, manual ROI drawing can demand much more time, depending on the total number of imaged ROIs (e.g., ∼40 min for 500 ROIs). Obviously, the segmentation with a hexagonal grid is instantaneous, but it lacks single-neuron resolution.
11) Select the diameter (in μm) of each hexagonal ROI.
12) Draw the perimeter of a closed polygon mask that encloses the imaged optical plane region to be segmented in ROIs. Right-click inside the mask and select *Create mask* to define it. For the *Use hexagonal grid of ROIs* option, the obtained hexagonal ROIs inside the mask will be displayed and results will be saved.
13) Move the *Gamma* and *Contrast* sliders in the GUI to optimize the display of the image and better visualize the segmentation (i.e., it will not affect the performance of the segmentation algorithm for the *Automatically detect single-neuron ROIs* option). When satisfied with the image, click on *Continue*.
14) Fluorescent labeling may be uneven in the imaged optical plane. To correct for this, move the *Local contrast scale* slider to perform a local intensity normalization of the image. Labeled cells should be clearly visible (even in originally darker image regions) and intensity should not vary significantly across the imaged plane. When satisfied, click on *Done*.

##### CRITICAL

It is this intensity-normalized image that will be segmented. If in the subsequent steps you are not able to obtain a satisfactory ROI segmentation, you will always be able to return to this step and perform a new normalization with a different value for the *Local contrast scale* by clicking on *Adjust gamma again* on the GUI.
15) If the fluorescent reporter labels nuclei, click on *Labeled Nuclei*, otherwise on *Unlabeled Nuclei* (the currently selected option will be indicated in red). This option is useful when using GCaMP indicators which usually do not enter the nucleus, in contrast to synthetic dyes (e.g. OGB-1 AM) and GCaMP NLS vectors.
16) Move the *Border thresh* and *Cell center thresh* sliders to adjust *thr_Neuropil_* and *thr_soma_*, respectively. Cell somata should be marked in red, and neuropil regions in blue (see left panel in Fig. 6a). Click on *Find ROIs* to inspect the preliminary automatic segmentation obtained using the selected parameters.

##### CRITICAL

Avoid merging neighboring cells by setting *thr_soma_* too high or over-segmenting cells by setting it too low. Labeling the neuropil in blue with an appropriate *thr_Neuropil_* will instruct the algorithm to avoid looking for ROIs in these regions. Fine-tuning of this parameter is not important when cells are tightly packed (e.g., Fig. 6), but will be of great help when cells are sparsely distributed (e.g., Fig. 2a,b).
17) Preliminary ROI perimeters will be shown in red. If they do not enclose cells completely, click on *Bigger ROIs*. If they exceed the cellular areas, click on *Smaller ROIs* (the currently selected option will be indicated in red). If results are not satisfactory, return to step 16. Otherwise, click on *Done*. You will always be able to return to steps 16-17 by clicking on *Find regional minima again* on the GUI.
18) Interactively impose several morphological criteria (ROIs’ minimal and maximal areas, intensities and circularities) to filter undesired ROIs (Fig. 6b). For this, move the corresponding sliders to any desired value, and click on *Run* in order to see the filtering effect. When finished, click on *Done*. You will always be able to return to this step by clicking on *Find automatic ROIs again* on the GUI.
19) Curate the ROI segmentation obtained (for the *Automatically detect single-neuron ROIs* option; Fig. 6c); define ROIs (for the *Manually draw all ROIs* option):
  i) To add new ROIs, click on *Draw ROIs*, click along the perimeter of every new ROI, and close the perimeter with a final right-click. Repeat this last procedure to add as many ROIs as desired. When done adding new ROIs, click on *Stop Draw ROIs* and draw an additional ROI. This last ROI will not be taken into account, but all the previously drawn ROIs will be defined.
  ii) To select multiple ROIs for deletion, click on *Delete ROIs* and right-click once on each ROI to delete, then click on *Stop Delete ROIs* and right-click once more in any place on the image. Again, this last selection will have no effect, but all the previously selected ROIs will be deleted.
  iii) In order to delete multiple ROIs in a specific region, select *Delete ROIs Area*, draw the perimeter of a closed polygon mask that encloses the concerned ROIs, right-click inside the mask and select *Create mask* to remove all ROIs inside the mask.
  To better visualize the imaged optical plane, click on *Hide ROIs* to stop displaying ROI perimeters for a few seconds. When finished, click on *Done* and results will be saved.

### Computation of ROIs’ relative fluorescence variations (∆F/F0) and detection of significant transients

20) Run the script *ProcessFluorescenceTraces.m*. The goal of this script is to determine the significant fluorescence transients in the data. Select the file with the *_ALL_CELLS.mat* suffix which contains the ROIs’ fluorescence traces.
21) Set the imaging parameters: the sampling frequency of the imaging video, and the fluorescence decay time constant of the reporter (τ; default value is the one reported for GCaMP3^66^, but for the correct detection of significant fluorescence transients, the τ of the imaged calcium reporter should be used).
22) Set the parameters for a data sanity test to eliminate ROIs whose fluorescence dynamics displays artifacts or have low signal-to-noise ratios (SNRs). The parameters are: the minimal number of pixels per ROI; the minimal ROI fluorescence relative to the baseline, used to detect ROIs whose fluorescence is too dim (lower parameter values will require more fluorescence from adequate ROIs); and the maximal sudden decrease in ROI baseline fluorescence, which detects ROIs whose baseline fluorescence is not stable (higher parameter values will require more stable fluorescence from adequate ROIs). ROIs that do not satisfy these criteria will be displayed, removed from the *_ALL_CELLS.mat* file, and will not be further analyzed (a backup of the original data will be saved with an *_ALL_CELLS_Original.mat* suffix).

##### CRITICAL

This step is intended to remove ROIs that could compromise the automated analysis. In our experience, the suggested default parameters are sufficient to assure data robustness, and they do not eliminate adequate ROIs. However, if the script fails to proceed, modify these values to increase the constraints on ROI data.
23) Choose if you want to subtract the local neuropil signal from each ROI fluorescence time series. If you answer *Yes*, you will be able to set the parameter α for the subtraction.
24) Select the method for the estimation of the ROIs’ baseline fluorescence. If you select *Average fluorescence on time window*, you will set a time window over which a constant basal fluorescence *F0* will be calculated. If you select *Smooth slow dynamics, F0* will be set equal to *F_smooth_*, which tracks the slow variations of the ROIs’ basal fluorescence level.
25) Select the method for the estimation of the noise in the signal (the characteristic scale σ of the ROIs’ baseline fluorescence noise). If you select *Gaussian model, σ* will be obtained by fitting a Gaussian model to the baseline fluorescence variations (recommended option). Otherwise, select *Standard deviation* and σ will be estimated as the standard deviation of the baseline fluorescence variations.

##### CRITICAL

The *Standard deviation* option should only be used if the imaged videos are short in time (i.e., shorter than ∼1000 imaging frames).
26) Select the method for the estimation of significant fluorescence transients in the data. You can either select *Static threshold* to use a constant fluorescence threshold (depending only on the ROI’s σ) or *Dynamic threshold* for one that is dynamic (using both ROI’s σ and the reporter’s fluorescence decay time constant τ to analyze fluorescence transitions across imaging frames). Alternatively, select *Import data* to import significant transients calculated using an alternative method. If you select the *Static threshold*, skip steps 28-29. If you choose the *Dynamic threshold*, skip steps 27 and 29. Finally, for *Import data*, skip steps 27-28.

##### CRITICAL

The *Static threshold* option is a simpler framework, but may be prone to detecting false positives in noisy data. The *Dynamic threshold* option is more computationally demanding, imposing larger constraints on the data (inferring data-driven noise models, hence making stronger assumptions on the data). However, it is more robust in the detection of real fluorescence transients in noisy data, especially for long videos (longer than ∼3000 imaging frames). Therefore, the choice depends on the SNR of the dataset, and the user’s outlook as to the analytical framework. We recommend testing both options and evaluating the result displayed at step 30.
27) Choose the minimal baseline-noise-scaled *∆F/F0* for an ROI’s significant fluorescence transient *(∆F/F0* x 1/σ; a measure of the SNR of the ROI’s transient). The default value for this parameter is 3 (i.e., only variations bigger than 3 times the baseline fluorescence noise level are considered significant). This parameter can be adjusted according to the SNR of the imaging video. The result of this section’s analysis is saved in a *Matlab* file with a *_RASTER.mat* suffix.
28) Set the minimal confidence (based on the noise model) that an ROI fluorescence transient is not noise (default is 95%). These values can be adjusted according to the SNR of the imaging video. The result of this section’s analysis is saved in a *Matlab* file with a *_RASTER.mat* suffix.
29) Select a *.mat* file containing a *T* × *N* array called *significantTransients*, where *T* and *N* are the number of imaging frames and ROIs, respectively. This array should contain a 1 at position (*i, j*) if the ROI *i* at imaging frame *j* showed a significant fluorescence transient, or 0 otherwise. The result of this section’s analysis is saved in a *Matlab* file with a *_RASTER.mat* suffix.
30) Select if you want to display examples of ROI *∆F/F0* traces along with the significant transients obtained highlighted in red (Figs. 2 and 6d). If you select *Yes*, you will then set the number of example traces to plot (ordered by level of ROI activity), and the *∆F/F0* scale of the plot. Once plotted, you will be asked if more traces should be displayed.

### Analysis of neuronal responses

31) Run *RunResponseMapping.m* and select the file with the *_RASTER.mat* suffix, produced either through this toolbox, or with data imported from other methods (for the format of this file, see *Imported Fluorescence Data File* in Box 2).
32) Select the *.mat* file with the timing information of the experimental events to be associated with ROI responses (for the format of this file, see *Response Analysis Input File* in Box 2).
33) Set the parameters for the procedure: the duration of each experimental event, and the duration of the time window, locked to event onset, over which ROI responses will be calculated. A graph will display the spatial topography of ROI responses (Fig. 4a).
34) To improve visualization of the data, click on *Remap colors* to adjust the *saturation* and *value* channels (if only one kind of experimental event was provided in step 32, color *saturation* will not be used). This will modify how response features are color scaled. Set the offset, lower and upper bound of the channels. Both the original and rescaled channels will be displayed (Fig. 4b), along with the spatial topography of ROI responses using the new color code (Fig. 4c).
35) Add or remove a transparency to the drawn ROIs proportional to their corresponding *value* channel by clicking on *Transp. on/off* (default is transparency on).
36) To display the average and single-trial responses and the tuning curve of a particular ROI (Figs. 4d,e), click on *Select ROI* and right-click on the ROI of interest. Change the layout of the average and single-trial responses by clicking on *Plot order* and choosing the number of rows used to display them.
37) When finished, click on *Save* and the calculated responses will be saved in a file with a *_RESPONSE_MAP.mat* suffix.

### Detection of neuronal assemblies

38) If you wish to find assemblies for a multi-plane volumetric imaging experiment, run *CombineMultiPlane.m* and select all the *_RASTER.mat* suffix files that correspond to each imaged plane (for data imported from other methods see *Imported Fluorescence Data File* in Box 2). This will generate files with *_MultiPlane_ALL_CELLS.mat* and *_MultiPlane_RASTER.mat* suffixes that you should use in the next step. Otherwise, for single-plane imaging experiments, skip this step.
39) Run the program *FindAssemblies.m*, and select the file with the *_RASTER.mat* suffix (for data imported from other methods see *Imported Fluorescence Data File* in Box 2).
40) Select the clustering algorithm to be used. For *PCA-promax*, skip steps 44-45; for *k-means*, skip steps 41-43 and step 45; and for *hierarchical clustering*, skip steps 41-43.
41) Select a cut-off value for the parameter that determines if a neuron belongs to an assembly (i.e., the z-scored maximal loading on the rPCs, *zMax)*. If you choose *No, zMax* will be set to the default value of 2; skip step 42. Otherwise, select *Yes* and continue with the next step.
42) A graph will display the probability density of *zMax* of all ROIs. This distribution should be multimodal, where ROIs that do not significantly correlate with other ROIs are concentrated in the first distribution peak. Select the *zMax* value that best separates this first peak from the rest of the distribution (see Fig. 7a). For better visualization you can optimize the smoothing level of the distribution by moving the *Smooth parameter slider* on the right of the graph and clicking on *Test smoothing* to inspect the distribution. To choose the value of *zMax*, click on *Select cut-off* and then click on the corresponding distribution point.
43) A window will display two population activity histograms and a raster plot of neuronal activity. The red line (determined by the *zMax)* on the raster plot separates neurons that belong to assemblies (above the line) from those that do not (below the line). Neurons above this line should participate in episodes of synchronous activity of distinct neuronal subpopulations (i.e., showing correlated activities). The middle population activity histogram corresponds to these assembly neurons. The population activity histogram of neurons below the red line is shown in the bottom panel. The activities of these latter neurons should look relatively uncorrelated with the rest of the neurons. Explore the raster plot by using the *Pan* and the *Zoom* icons in the figure’s toolbar. Press *Enter* if you are satisfied with the threshold selected. If not, answer *No* and you will be redirected to the *zMax* threshold selection of the previous step.

##### BOX 2 Imported ROIs

For an imaged plane of *P_x_* rows and *P_y_* columns of pixels, a *.mat* file containing a variable called *importedROIs* storing a logical binary *P_x_* × *P_y_* matrix that represents a mask of all desired ROIs pooled together, where a value of 1 marks ROI pixels and a value of 0 pixels outside ROIs.

###### Imported Fluorescence Data File

For *T* imaging frames and *N* ROIs, a *.mat* file with the *_RASTER.mat* suffix contains the following variables:

*deltaFoF:* a T × N matrix of the ΔF/F0 time series of all ROIs.
*raster:* a *T* × *N* matrix. For each column (i.e., each ROI), it is filled either with zeros for frames with non-significant fluorescent transients or with the *∆F/F0* values of the frames with significant transients. If the user does not wish to plot significant trial responses in the response analysis module, or if all the *∆F/F0* values should be considered in the module for the detection of the assemblies (instead of only the significant transients), it should be a *T* × *N* matrix filled with ones.
*movements: T* × 1 binary array, with ones for frames where an imaging artifact was found, and otherwise zeros.
*dataAllCells.avg:* average image of the imaging file, showing the anatomy of the imaged plane.
*dataAllCells.cell_per:* an *N*×1 cell array, containing the perimeter coordinates for each ROI.
*dataAllCells.cell:* a 1 × *N* cell array, containing the pixel indexes of each ROI.

###### Response Analysis Input File

A *.mat* file containing a 1 × *S Matlab cell array* called *mapData*, where *S* is the total number of experimental event types (for the OT case study, these are the azimuth positions of the visual stimuli). Each layer of the *cell array* (e.g., each stimulation position), should contain the following variables:

*label:* a string with the corresponding event label for the *Mapping parameter* color bar (e.g., “−35^°^”).
*value:* a scalar with the corresponding parametric value of the mapped event (e.g., “35”).
*onsetTime:* a vector of scalars corresponding to the timing information in seconds of each event trial (e.g., “[10.0 25.2 37.1]” for 3 trials).

###### Time Tags for Assemblies File

A *.mat* file containing an *N* × 1 vector called *timeTags*, where *N* is the total number of imaging frames. In this vector, frames with no particular tags are labeled with a value of 0, and those associated with a given tag are labeled with the corresponding tag number, according to your experimental setup (for example, frames tagged with a number 1 for when the animal is running, a number 2 when there was an optogenetic stimulation, a number 3 when a visual stimulation was presented, etc.).

###### File With ROI Features to Test Against Surrogate Assemblies

For evaluating certain features of the assemblies, the user must provide a *.mat* file with the features to be tested against the control surrogate datasets. To test *F* features, this file should contain a *Matlab cell array* called *variable*, whose dimension should be *F*×1 (that is, one *cell* for each feature). For a total population of *N* neurons, the dimension of each *cell* in *variable* depends on the kind of feature to be tested, as described below.

Single-neuron features: for a scalar feature *F_i_* (e.g., the average activation rate of each neuron), *variable{F_i_}* should have a dimension of *N*×1. For a vectorial feature *F_i_* sampled at *m* conditions (e.g., the response tuning curves of neurons with respect to a visual stimulus at *m* positions of the visual field), *variable{F_i_}* should be a *Matlab structure*. In this structure, the values of the features are stored in a Nxm array called *variable{F}.y* (for the tuning curves example, this corresponds to the *ΔF/F0* neuronal responses), and a 1xm array called *variable{F}.x* must store the condition values (the tested visual field positions).

Neuronal-pair features: for a pair-wise scalar feature *F_i_* (e.g., the pair-wise activity correlations), *variable{F_i_}* should have a dimension of *N×N*.
44) Select if the dimensionality of the data should be reduced through PCA, the total number of clusters to look for in the data, and the distance metric to be used *(euclidean* or *correlation)*.
45) Select the linkage algorithm to calculate the hierarchical tree: *single* or *complete*.
46) A series of figures will display the spatial topographies of all the assemblies found in the data. Results will be saved in a Matlab file with a *_CLUSTERS.mat* suffix.

##### TROUBLESHOOTING

To evaluate the validity of the obtained assemblies, run steps 38-47 choosing different methods and parameters, and then run the program *indexesAssemblies.m* on all the *_CLUSTERS.mat* files obtained. This program will test the clusterings in these files, by calculating and displaying the *Davies-Bouldin*^*69*^(DB) and the *normalized Hubert’s Γ*^*70*^ cluster quality indexes. You should choose the *_CLUSTERS.mat* that results in the lowest DB and the highest Γ indexes. Furthermore, the relevance of the assemblies obtained can also be qualitatively assessed by inspection of their anatomical topography in relation to experimental events (e.g., sensory or motor functional maps), especially when imaging large brain regions.
47) Run *AssembliesActivations.m*. This script will analyze all the population activity bouts and estimate if they can be confidently assigned to specific neuronal assemblies. Select the file with the *_CLUSTERS.mat* suffix obtained in the previous step. Select the threshold *p*-value to calculate the significance of the activation of an assembly (default is 0.01, lower values correspond to a higher statistical confidence). Results will be saved in the *_CLUSTERS.mat* file.

### Exploratory analysis of assemblies

48) Run the *AssembliesTopographicalOrganization.m* script to evaluate and display all the assemblies color-coded according to their spatial or temporal organization (the obtained organization data will be saved in a *_ORDER_TOPO,mat* suffix file). Select the file with the *_CLUSTERS.mat* suffix and choose if assemblies should be organized according to the similarity of their activity dynamics (click on *Similarity of dynamics*) or according to their spatial distribution along a particular anatomical axis (choose *Along topographical axis*). For the former temporal option, the color-coded assemblies will be displayed (e.g., Fig. 5), and the image will be saved in a file with a *_assemblies_topo_temporal.png* suffix. For the latter spatial option, continue to next step.
49) Assemblies will be organized (and color-coded) according to the spatial projection of their topographical centroids over a user-defined axis. This axis can be drawn (click on *Along curve*, and continue to step 50). Alternatively, it can be automatically set as the left-to-right axis (click on *Left-right)* or the bottom-to-top axis (*Bottom-top* option). For these two latter options, skip step 50, and the color-coded assemblies will be displayed (saved in a file with a *_assemblies_topo_spatial.png* suffix).
50) The imaged plane may contain multiple brain regions with their corresponding topographical axes. Set the number of regions. For each region, draw a polygon mask that encloses the region, and finish the mask by right-clicking inside the drawn polygon and selecting *Create mask*. For each region, define the corresponding axis by drawing a curve with several mouse clicks, finishing it with a final right-click. The axis curve will be displayed with a color code along its length (Fig. 7c). If you are not satisfied with the curve, click on *No* to the *Satisfied with the curve?* question and draw it again. Once finished, the color-coded assemblies will be displayed (Fig. 7d; saved in a file with a *_assemblies_topo_spatial.png* suffix). Assemblies will be colored according to the selected axis: each assembly will be displayed with the color of the axis point that is physically closest to the assembly’s centroid.
51) The script *AssembliesDynamicsVisualization.m* allows you to explore and evaluate the dynamics of the activation of the assemblies. You will also be able to incorporate multiple time tags corresponding to experimental events (e.g., behavioral epochs, sensory and/or neuronal stimulation times, etc.). Run it and select the *_CLUSTERS.mat* file. Select the *_ORDER_TOPO.ma*t file from the previous step. If you want to incorporate experimental time tags, click on *Yes* and select a *.mat* file where the tags are stored (see *Time Tags for Assemblies File* in Box 2 for the required file format).
52) A figure will display the assemblies’ activation dynamics in relation to the experimental time tags informed in the last step (Fig. 8). The buttons on the left allow the choice of which assemblies to display. For visualization of all the assemblies, click on *All*. For a selection of assemblies, click on *Selection* and provide a space-separated list of assemblies to be displayed. You can explore the raster plot of the top panel using the *Pan* and the *Zoom* icons in the figure toolbar, and the middle and bottom panel graphs will be displaced in synchrony with the top panel.

### Generation of control surrogate datasets for statistical analysis of assemblies’ features

53) Run the program *SurrogateData.m*. Select the corresponding *_CLUSTERS.mat* file. Choose if you wish to produce *Random surrogate assemblies, Topographical surrogate assemblies* or *Both* (see Box 1). Set the number of surrogate assemblies to produce per original assembly (default is 100). The surrogate assemblies will be saved in a file with a *_SURROGATE_CLUSTERS.mat* suffix.

##### TIMING

Depending on the total number of topographical surrogate assemblies, this program could perform calculations for a few minutes up to an hour.
54) The program *AssembliesVsSurrogate.m* will compare user-defined features of original and surrogate assemblies. For this, the program will pool single-neuron features (e.g., the average frequency and/or intensity of activation, the neuronal tuning curve, the neuronal identity, etc) and/or neuronal-pair features (e.g., the pair-wise activity correlation, the pair-wise topographical distance, etc.) according to the neuronal groups represented in the original and surrogate assemblies. You can test *F* features by providing an appropriate *.mat* file (see *File with ROI Features to Test Against Surrogate Assemblies* in Box 2 for the required file format). Run *AssembliesVsSurrogate.m*, select the *_CLUSTERS.mat* file, then the *.mat* file for the neuronal features, and the program will compare against the surrogate assemblies stored in the *_SURROGATE_CLUSTERS.mat* file created in the previous step.
55) A set of graphs will be displayed for each kind of surrogate assembly used *(Random*, or *Topographical;* Fig. 9a). For scalar variables, each graph will display a histogram of the pooled features of the original (top panel) and the surrogate assemblies (bottom panel; Fig. 9b,c). For vectorial variables, histograms are replaced by a graph of the average feature curve (black line) and pooled single-neuron curves (gray lines). The final pooled features will be saved in a file with a *_ASSEMBLIES_vs_SURROGATE.mat* suffix that contains the original and surrogate assemblies’ features in Fx1 *cell arrays* named *varAssemblies* and *varNMs*, respectively. Use any pertinent statistical test of your choice to assess the significance of the original assemblies’ features when compared against the surrogate assemblies.

## RESULTS

The file called *OT_reg_smooth.tif* (in the folder *Test data* of the toolbox distribution) contains data from a two-photon imaging experiment of the optic tectum (OT) of transgenic GCaMP3-expressing zebrafish larvae. It will be used as a case study in this section. In this experiment, the OT was imaged while periodically stimulating the larva with single light spots in different positions of the larva’s visual field to map the OT functional retinotopicity. With this case study, we wish to illustrate the use of the toolbox and, furthermore, assist users who wish to adapt this procedure to their own datasets. We also comment on the results obtained when analyzing other kinds of data.

We will focus on the more involved procedures of the protocol, skipping some of the simpler initial preprocessing steps. Hence, this case study has been already registered for drifts of the optical plane using *SmoothRegistration.m* as explained in step 1 (for a demonstration of the registration effect, compare with the unregistered *OT.tif* file), and movement artifacts have already been detected as explained in steps 2-9 (artifacts were detected with default parameter values, and were stored in *OT_reg_smooth_ARTIFACTS.-mat)*. The user is encouraged to perform all the protocol steps on this case study, but we also provide all the remaining *Matlab* files that we obtained with the tutorial (see folder *Test data\Processed files)*, so that users can compare their own results.

For the OT case study, Figures 6a-c show the parameters selected for the intermediate steps and a typical result of using the semi-automatic ROI detection algorithm, as described in steps 12-19. Figure 2 illustrates the applicability efficiency of the algorithm for the analysis of markedly different imaging data. The identification of ROIs in the OT case study is rather difficult, due to the high density of neuronal somata and the thin GCaMP3-expressing boundaries between neurons. Nevertheless, with the chosen parameters, it can be completed in ∼10 mins (depending on user experience), obtaining 854 ROIs. For both mouse cortical datasets (Fig. 2a,b) it took less than 5 mins to obtain 149 and 505 ROIs, respectively. For the large-field dataset in zebrafish (2178 ROIs), it took ∼10 mins (Fig. 2c). Finally, for the light-sheet imaging dataset we obtained 438 ROIs in less than 10 mins (Fig. 2d). Typically, the user may need to curate 5-15% of the ROIs automatically obtained, depending on imaging quality. For the present examples, after setting the parameters for ROI segmentation (steps 12-18) and filtering (step 19), we had to curate 9% of the ROIs for the OT (Fig. 6) and both mouse cortical datasets (Fig. 2a,b), while for the large-field zebrafish dataset, 8% of the ROIs were curated (Fig. 2c). For the single-plane single-photon light-sheet imaging dataset shown in Figure 2d, we had to curate 15% of the ROIs, due to the lower spatial resolution of this technique. Since simultaneous multi-plane light-sheet imaging does not guarantee single-neuron resolution, we used an array of hexagonal ROIs (Fig. 5; steps 11-12). Manual curation of ROI segmentation is the protocol step that demands the most user intervention.

We continue with the inference of significant *∆F/F0* transients, which can also be applied to a variety of imaging conditions (Figs. 2 and 6d). For the present zebrafish OT case study and the large-field zebrafish dataset, the data sanity test was run with default parameters (step 22), no neuropil signal correction was used (step 23), the ROIs’ baseline noise σ can be estimated with the *Gaussian model* (step 25), and the significance of fluorescence transients can be inferred with the *Dynamic threshold* method with default parameters (steps 26-28). The mouse cortical data was similarly analyzed, but with neuropil signal correction, and the appropriate τ of the calcium reporters. For the light-sheet imaging datasets in Figs. 2d and 5, σ was also estimated with the *Gaussian model*, but due to the large size of the datasets, significant transients were obtained with the faster *Static threshold* method (step 26). Nevertheless, we encourage the user to experiment with both changing parameters and implementing the alternative procedures of the protocol (e.g., estimation of σ through *Standard deviation*, or importing data from other methods in step 29).

To illustrate the capabilities of the response analysis module (steps 31-37), we present the results obtained for the OT case study (Fig. 4). Here, stimulation consisted of 3 trials of 4^°^ light spots presented at different visual-field azimuth positions (−45° to 45^°^ in 5^°^ steps, 0^°^ represents directly in front of the larva). The file with trial stimulation information used by the response analysis module (step 32) is *OT_reg_s-mooth_TimeTagsMapping.mat*. Despite the small number of trials analyzed (3 per azimuth angle), Figure 4 demonstrates the potential of mapping the neuronal preferred stimulus, selectivity and response strength with a *hue-saturation-value* (HSV) color code, which allows an efficient visual inspection of the topography neuronal responses (compare the information conveyed by mapping in Fig. 4c with that of Supplementary Fig. 2b, where the *saturation* and *value* channels were rendered flat). However, since the range of neuronal responses can be quite large, the HSV mapping typically must be optimized to avoid obscuring/smothering weaker but significant responses. The module’s GUI greatly facilitates this optimization (Fig. 4b) by revealing relevant biological information (Fig. 4c) that would otherwise be obscured (Fig. 4a). Furthermore, the responses of any ROI of interest can be interactively explored (Fig. 4d). Importantly, the versatility of this module design allows the study of the neuronal coding of any other kind of (parametric) experimental variable (e.g., stimulation, behavioral) whose time tags are known, in an automatic manner.

For the detection of neuronal assemblies in the OT case study, we used the *PCA-promax* method. We chose to manually select the single input parameter required by the program, the *zMax* threshold (step 42, Fig. 7a). Examples of the spatial topographies of the detected assemblies are shown in Figure 7b, which reveal a majority of spatially compact neuronal clusters. This is an expected result given the light-spot visual stimulation used and the retinotopic organization of the OT, where neighboring positions in the visual field are mapped on neighboring positions in the OT^67,68^. In Figure 5 and **Supplementary Videos 1** and **2**, we demonstrate the result of using the *PCA-promax* method to detect assemblies in large-scale multi-plane light-sheet imaging (i.e., spontaneous activity of >40,000 simultaneously monitored ROIs scanned at 2.1 Hz in a zebrafish larva). Several bilaterally symmetric assemblies and unilateral assemblies with their corresponding contralateral counterparts were found. This illustrates the richness of the biological information obtainable with this clustering algorithm, which surpasses other popular algorithms *(k-means)* in exposing the spatio-temporal organization of this high dimensional dataset (compare Figs. 5g and 5h). We encourage users to detect assemblies in the OT case study using the *k-means* and *hierarchical* clustering algorithms implemented in the module, and experiment on the effect of changing the clustering options (e.g., distance metric, linkage option). The validity of the assemblies obtained with the different methods can be evaluated through the provided cluster quality indexes (see Troubleshooting of step 46).

The retinotopic-like organization of the observed assemblies in the OT case study can be further confirmed with the exploratory analysis module, without any information about the visual stimulation events (Figs. 7c,d; steps 48-50). This demonstrates the potential of the assemblies’ module to uncover latent organizing principles of the data in an unsupervised manner. The assemblies’ activation dynamics can also be further studied using steps 51-52. In Figure 8, we present an example of the dynamics of a few selected assemblies, confirming that most of them are engaged by some light-spot stimulations, as indicated by the experimental time tags provided in the *OT_reg_smooth_TimeTags.mat* file. Finally, assemblies’ features can be compared against control surrogate assemblies (Fig. 9; steps 53-55). In this case study, as a feature input file for step 54, we used the *OT_reg_smooth_VARIABLE.mat* file. This file stores the average ROI activity levels in the vector *variable #1*, and the ROI-ROI activity correlations in the matrix *variable #2*. We found that compared to *Topographical surrogate assemblies*, assemblies do not show any particular trend in the average activity level of their component ROIs (Fig. 9b), but they do significantly group correlated ROIs (Fig. 9c). The same trend was observed when comparing with *Random surrogate assemblies*. This example allows the user to further validate the clustering results, which effectively groups correlated ROIs, without showing a bias for the most active ones (a common concern in clustering procedures). Nevertheless, this module is not limited to methodological questions such as these. It aims to evaluate scientific questions, such as assessing whether assemblies group ROIs according to their functional properties (such as those obtainable with the response analysis module), quantified by ROI features that the user provides in a feature input file. Furthermore, cross-talk between this module and the results revealed by the exploratory analysis module (see Fig. 1) allows the user to properly choose relevant features to be tested.

## ACKNOWLEDGEMENTS

We thank Simon Peron, Karl Svoboda and collaborators, and also Nicholas Priebe, Boris Zemelman and collaborators for sharing with us the mouse somatosensory and visual cortex data, respectively, through the *crcns.org* data sharing portal (datasets ssc-1 and pvc-10). We thank the members of the Sumbre group for helpful discussions and continuing testing of the toolbox. We thank the Marin Burgin lab at IBioBA for their support. This work was supported by EraSysBio+ Zebrain, ERC stg 243106, ANR-10-LABX-54 MEMO LIFE, ANR-11-IDEX-0001-02 PSL* Research University, Avenir grant INSERM, Argentine Agency from the Promotion of Science and Technology (PICT 2013-0182), and Structural Convergence Fund for MERCOSUR (FOCEM).

## AUTHOR CONTRIBUTIONS

S.A.R. designed the toolbox, developed the software, performed imaging experiments and analyzed datasets. V.P.-S. developed software and interactive user interfaces. A.J. and A.C. contributed and analyzed light-sheet imaging data. J.B-W. Developed the *HuC:GCaMP5* zebrafish line. S.A.R. and G.S. wrote the manuscript.

---

**Supplementary Figure 1.**
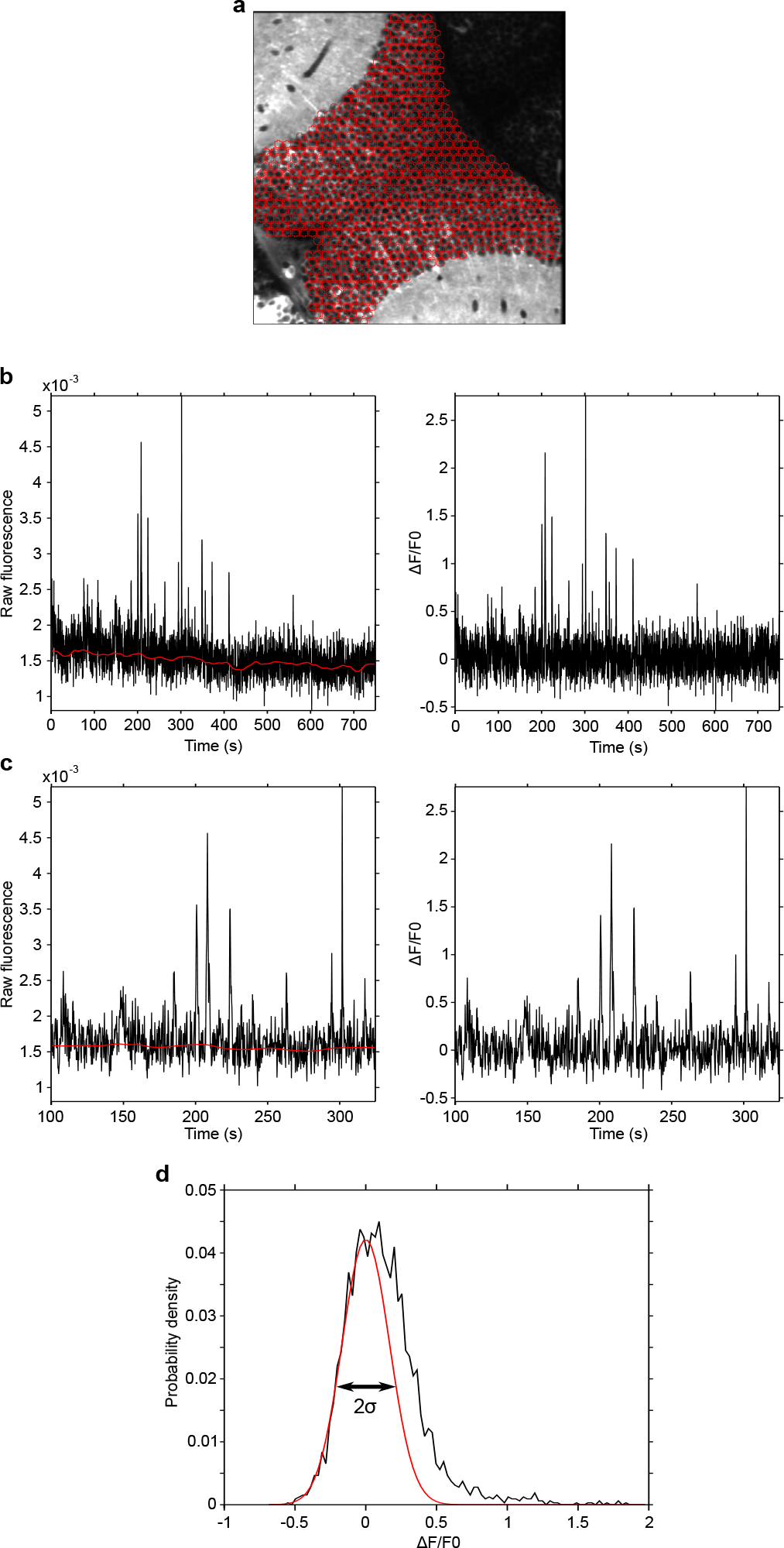
Pre-processing of fluorescence dynamics. **(a)** Red, hexagonal grid of 6 μm, over an imaged optical section of OT of a zebrafish larva pan-neuronally expressing GCaMP3. The region covered by the grid is defined with a user-drawn mask (step 12). (**b**) Left, raw fluorescence of a ROI (black) and the estimated *F_smooth_*. Right, *∆F/F0* obtained using *F_smooth_* as *F0*. Note how slow fluctuations are removed, producing a stable *∆F/F0*. (**c**) Zoom of **a.** Note how follows slow fluorescence variations, ignoring the fast, neuronal activity related fluorescence transients. (**d**) Estimation of the baseline fluorescence noise (**σ**) of a ROI. Black, normalized histogram of the ROI’s *∆F/F0;* red, Gaussian fit to the negative fluorescence *∆F/F0*.

**Supplementary Figure 2.**
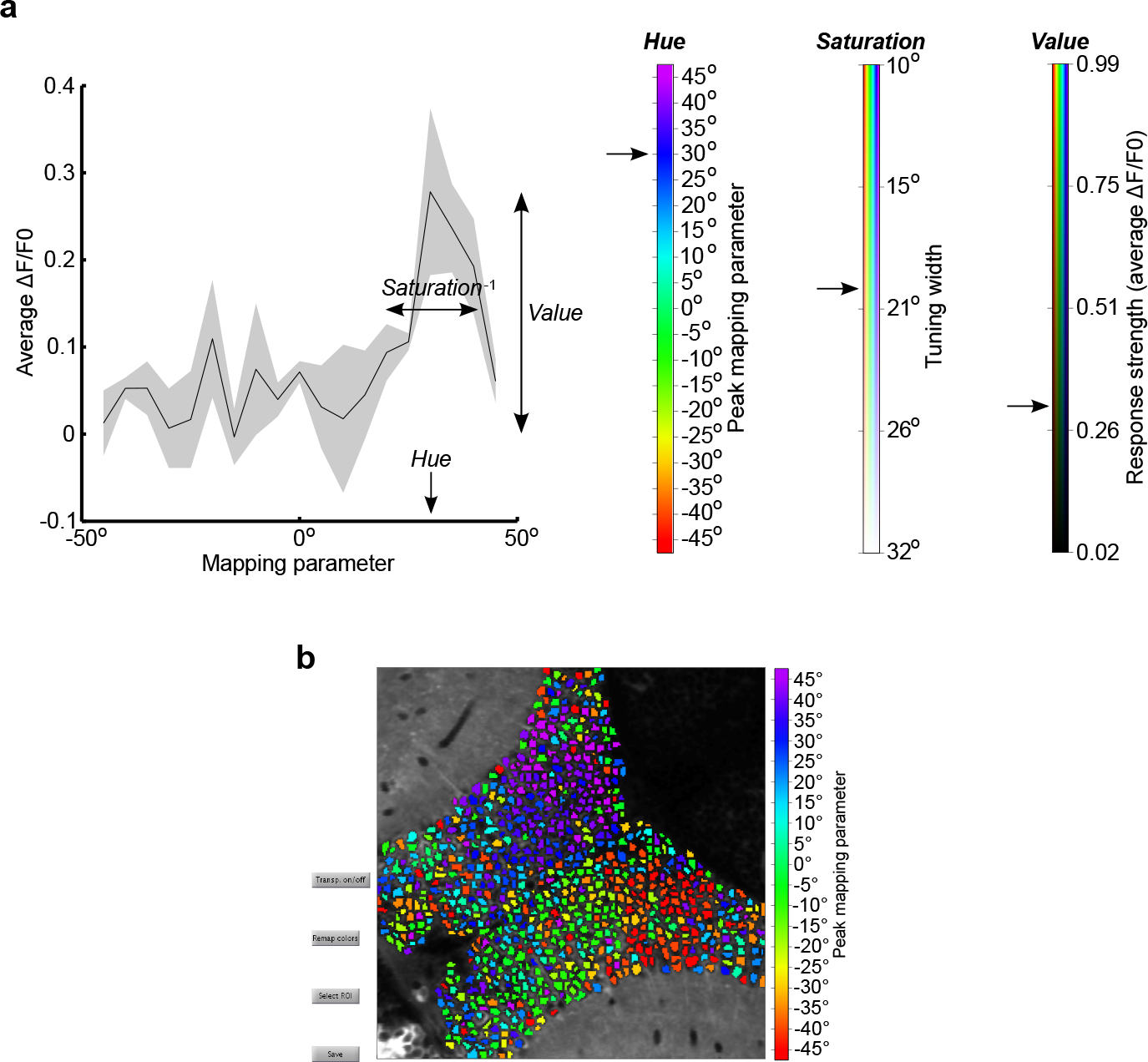
Mapping of responses to *HSV* color-code. **(a)** Schema illustrating the definition of the hue, *saturation* and *value* to visually represent ROI responses in a color-code. Left, ROI tuning curve. Arrows indicate the particular hue, *saturation* and *value* of this ROI. Right, colorbars representing the range of hue, *saturation* and *value* for all the imaged ROIs. Arrows indicate the color-code parameters for the ROI tuning curve shown in the left. (**b**) Display of neuronal responses only representing preferred stimulus (i.e., the peak mapping parameter). Same as Fig. 4, but disabling *saturation* and *value* channels, thus only representing preferred stimulus on the *hue* channel. The noisier image obtained underscores the utility of additionally representing the neuronal selectivity and response strength.

**Supplementary Video 1.** Volumetric distribution of example assemblies shown in Fig. 5a-f. Assemblies obtained with the *PCA-promax* algorithm. For each panel, the corresponding axial projection of the example assemblies is initially displayed, followed by their layout over each one of the 40 imaged optical planes (plane depths are indicated in lower right corners).

**Supplementary Video 2.** Volumetric distribution of all the assemblies shown in Fig. 5g. Assemblies obtained with the *PCA-promax* algorithm. The axial projection of all the assemblies found in the dataset is initially displayed, followed by each one of the 40 imaged optical planes (plane depths are indicated in lower right corners).

